# Competition within low-density bacterial populations as an unexpected factor regulating carbon decomposition in bulk soil

**DOI:** 10.1101/2020.11.16.384735

**Authors:** Alexandre Coche, Tristan Babey, Alain Rapaport, Laure Vieublé Gonod, Patricia Garnier, Naoise Nunan, Jean-Raynald de Dreuzy

## Abstract

Bacterial decomposition of organic matter in soils is generally believed to be mainly controlled by the access bacteria have to their substrate. The influence of bacterial traits on this control has, however, received little attention. Here, we develop a bioreactive transport model to screen the interactive impacts of dispersion and bacterial traits on mineralization. We compare the model results with two sets of previously performed cm-scale soil-core experiments in which the mineralization of the pesticide 2,4-D was measured under well-controlled initial distributions and transport conditions. Bacterial dispersion away from the initial substrate location induced a significant increase in 2,4-D mineralization, revealing the existence of a regulation of mineralization by the bacterial decomposer density, in addition to the dilution of substrate. This regulation of degradation by density becomes dominant for bacteria with an efficient uptake of substrate at low substrate concentrations (a common feature of oligotrophs). The model output suggests that the distance between bacteria adapted to oligotrophic environments is a stronger regulator of degradation than the distance between these bacteria and the substrate initial location. Such oligotrophs, commonly found in soils, compete with each other for substrate even at remarkably low population densities. The ratio-dependent Contois growth model, which includes a density regulation in the expression of the uptake efficiency, provide a more versatile representation than the substrate-dependent Monod model in these conditions. In view of their strong interactions, bioreactive and transport processes cannot be handled independently but should be integrated, in particular when reactive processes of interest are carried out by oligotrophs.

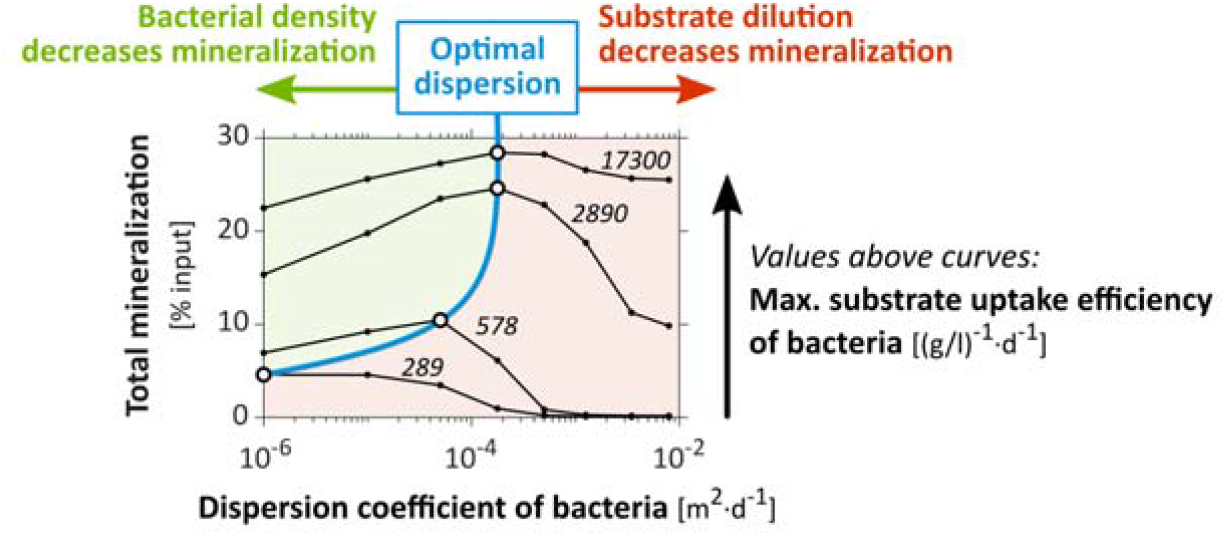

**Highlights:**

- The impact of spatial distributions on decomposition depends on bacterial traits
- Decomposition can be reduced by competition between bacteria even at low densities
- Bacterial density regulation counterbalances substrate accessibility regulation
- Regulation of decomposition by bacterial density is more acute for oligotrophs

## 1. Introduction

Organic carbon is involved in most ecological functions provided by soils (Bünemann et al., 2018). Its cycling in soil depends upon the activity of microorganisms. Soluble organic molecules are taken up as substrates by specific populations of soil bacteria, and degraded inside the cells by endoenzymes to provide carbon and energy. This is precisely the case for the 2,4-Dichlorophenoxyacetic acid (2,4-D) used in this study as a generic model compound (Don and Weightman, 1985; Pieper et al., 1988; Boivin et al., 2005). Bacterial degradation of soil carbon has generally been modeled with the Monod equation, where the specific substrate uptake rate is controlled by substrate concentration and bacterial traits such as the maximum specific growth rate, the yield (or carbon use efficiency) and the “maximum uptake efficiency” (e.g. Monod, 1949; Sinton et al., 1986; Cheyns et al., 2010). With the Monod equation, at the lowest substrate concentration, the specific uptake rate is linearly proportional to the substrate concentration. The proportionality factor is referred to here as the “maximum uptake efficiency” and it reflects the maximal ability of the cell to capture substrate molecules that collide with its membrane (Button, 1978, 1983). The maximum uptake efficiency can also be understood as the volume from which a cell can harvest substrate per unit of time, as used in some studies (Desmond-Le Quéméner and Bouchez, 2014; Nunan et al., 2020; Ugalde-Salas et al., 2020). Each bacterium is assumed to be exposed to the whole substrate concentration of its surroundings, without any limitation by the population density (Lobry and Harmand, 2006).

The direct contact (exposure) between bacteria and substrate depends on their spatial distributions (Holden and Firestone, 1997; Nunan et al., 2007). Bacteria and substrate are both heterogeneously distributed as a result of numerous biotic and abiotic processes (Dechesne et al., 2014; Kuzyakov and Blagodatskaya, 2015). There are complex feedback loops between these distributions, dispersive transport processes such as diffusion and hydrodynamic dispersion (Madsen and Alexander, 1982; Breitenbeck et al., 1988), and the bacterial activity itself such as consumption and growth (Poll et al., 2006).

Aggregated bacterial distributions, as observed at mm-scale for 2,4-D degraders (Vieublé Gonod et al., 2003), have been shown to decrease degradation rates when the distribution of substrate is homogeneous (Pallud et al., 2004; Dechesne et al., 2010). Yet, the role of bacterial metabolic traits on the impact of bacterial and substrate distributions on degradation remains mostly unknown, especially when substrate and bacteria are heterogeneously and dynamically redistributed in soils over µm-to-cm scales by numerous spatial disturbances (Madsen and Alexander, 1982; Breitenbeck et al., 1988; König et al., 2020). We investigated the extent to which bacterial activity and transport processes can be treated independently or should be integrated to characterize, understand and predict degradation under various advective, diffusive and dispersive conditions. The simultaneous characterization of the impacts of bacterial traits and transport parameters through their mutual interactions is methodologically challenging. It requires several well-controlled experiments in comparable degradation conditions, with specific spatial distributions of substrate and degraders in specific transport conditions, and a spatiotemporal monitoring of the different carbon pools.

Among the scarce relevant datasets (e.g. Dechesne et al., 2010), we used the two sets of cm-scale soil-core experiments performed by Pinheiro et al. (2015, 2018), in which the degradation of 2,4-D under different initial spatial distributions and transport conditions was measured in similar repacked soil columns. Mostly reported independently, they have shown first that the proximity between bacteria and the initial location of a heterogeneously distributed substrate exerts a strong control on mineralization. Mineralization was greater when bacteria were close to the initial location of substrate, even though most of the initial soluble substrate diffused away from its initial location. This was attributed to the fact that bacteria located far from the initial substrate location were only exposed to highly diluted substrate concentrations (Babey et al., 2017). However, the hydrodynamic dispersion of both bacteria and substrate away from their initial location caused a greater than four-fold increase in the mineralization of substrate that was not leached out, to the point that it almost reached the same performance as in homogeneous conditions in which there was no dilution (Pinheiro et al., 2018). The surprising increase in mineralization suggests a regulation of mineralization by population density compensating the effect of substrate dilution, the activity of bacteria being enhanced when their density is diluted by the dispersive percolation events. While such regulations by bacterial density have not yet been considered in soils, presumably because of the extremely low apparent bacterial densities found in soils (Young et al., 2008), they are well known in bioreactors, where they are usually modeled by the ratio-dependent Contois growth law (Contois, 1959; Harmand and Godon, 2007).

In order to determine the relevance of the putative bacterial decomposer density effect on decomposition, we developed a quantitative approach to model the two sets of experiments within the same unified framework (section 2). We assessed the relevance of previously developed models, improved the calibration of a Monod-based model and investigated an alternative Contois-based model (section 3). We discuss the implication of the results on the controlling factors of soil organic carbon cycling, on the relevant bacterial growth models and on the possible bacterial strategies (section 4).

## 2. Models and methods

### 2.1. Experiment scheme, geometry and initial distributions

We briefly introduce the experiments performed previously and highlight aspects of the experiments that are important for the modeling (**Fig. 1**). The full experimental setting is presented in the supplementary materials (**Fig. S1** and **Table S1**) for the sake of completeness. Soil columns were packed with two homogeneous or heterogeneous arrangements of soil cubes, either sterilized, or hosting the indigenous microbial communities (referred to as “degraders”) and amended with ^14^C-labelled 2,4-D (referred to as “substrate”). Two sets of experiments, referred to as “hydrostatic” and “percolation” conditions, were performed respectively with only substrate diffusion (Pinheiro et al., 2015), or with additional substrate and bacterial advection and dispersion caused by water percolation (Pinheiro et al., 2018). The initial locations of the bacteria and substrate were set in the model according to the experimental conditions (**Fig 1A**). Initial concentrations used in the model are detailed in Table 1. In the experiments, the mass of mineralized ^14^C derived from the degradation of the labelled 2,4-D was monitored at the core scale during at least two weeks (**Fig. 1B**). These data were used to confront the model processes with a physical system, as detailed in section 2.5.

**Table 1.**
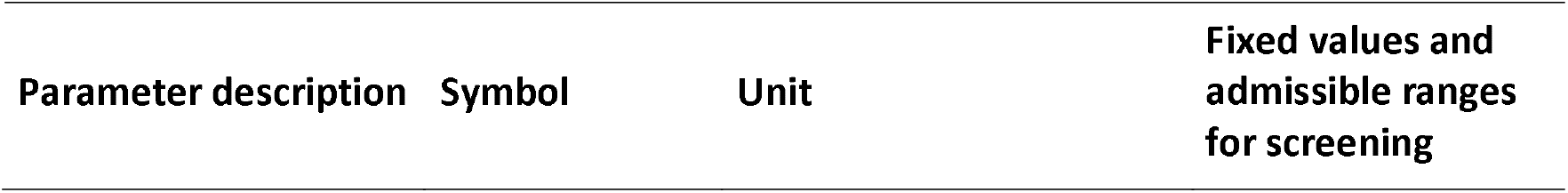
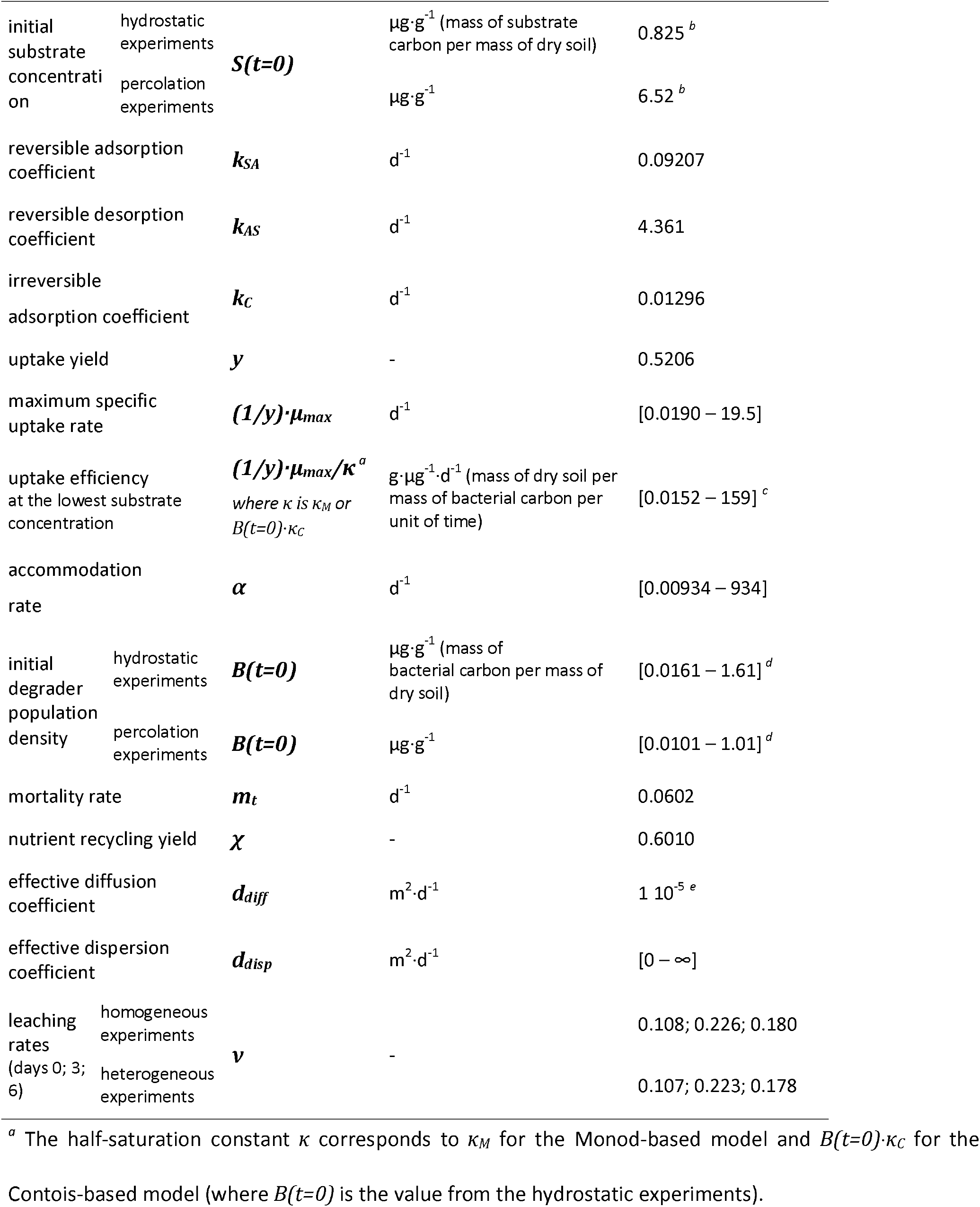
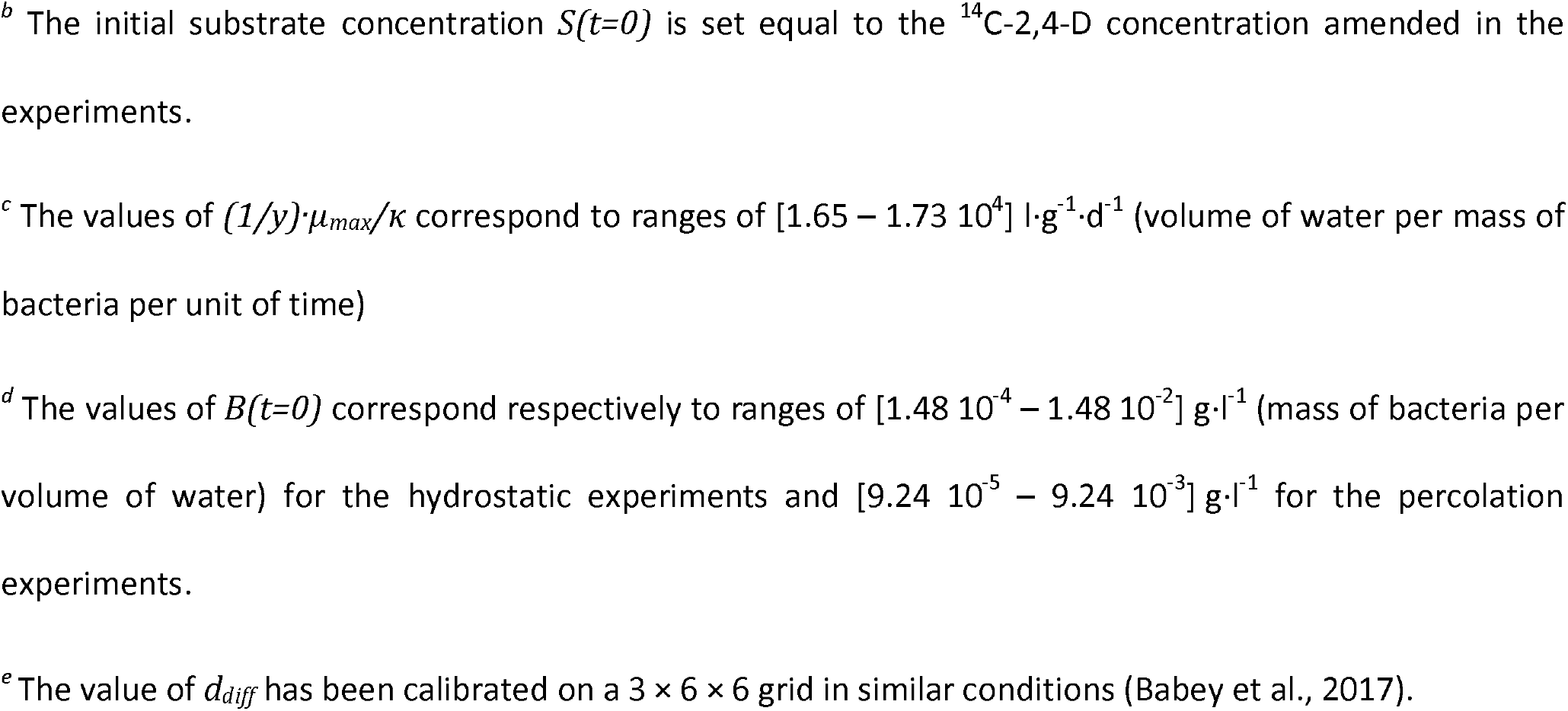
Values and range of values of the reactive transport model. The effective dispersion coefficient *d*_*disp*_ applies only to heterogeneous percolation experiments. *B(t=0)* is the initial density of bacteria in the natural cubes. It is considered 1.6 times smaller in the percolation experiments than in the hydrostatic experiments according to the initial experimental measurements.

**Fig. 1.**
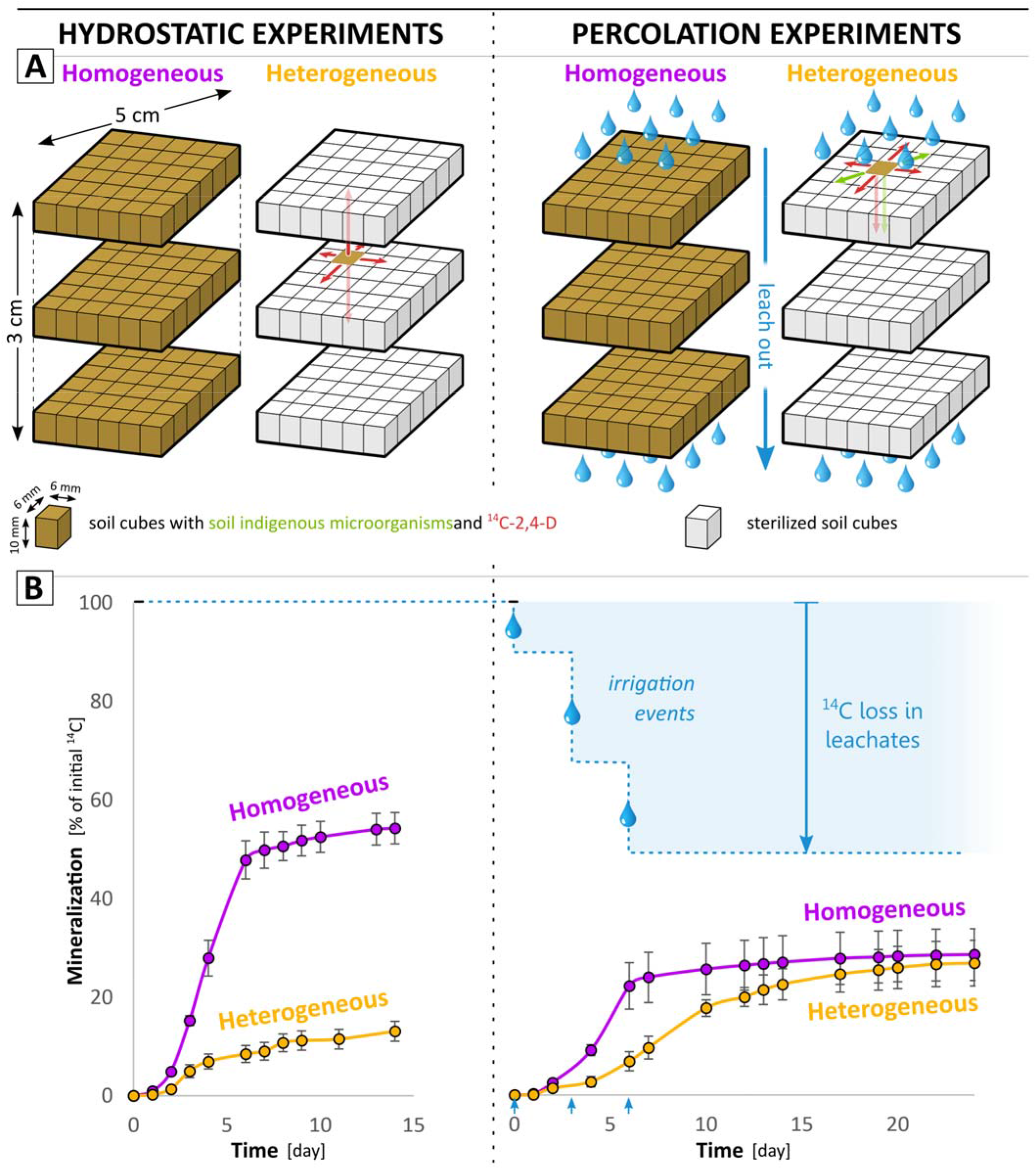
Model experimental design, geometry and initial distributions (**A**) based on previously performed experiments in hydrostatic (Pinheiro et al., 2015) and percolation (Pinheiro et al., 2018) conditions. The red and green arrows refer respectively to the 2,4-D and degrader modeled displacements. (**B**) Experimental cumulated production of CO_2_ (adapted from Pinheiro et al. (2018, 2015), permission for reproduction granted by Elsevier).

### 2.2. Bioreactive model

The bioreactive model extends the model published by Babey et al. (2017) (**Fig. 2**) to account for Contois growth law as an alternative to Monod’s. The sorption processes, the bacterial lag phase and the nutrient recycling described below were previously discussed and their use justified in Babey et al. (2017) to consistently represent the experimental data. The *r(·)* notation expresses the reaction rates of the biochemical dynamics that are expressed as follows:

**Fig. 2.**
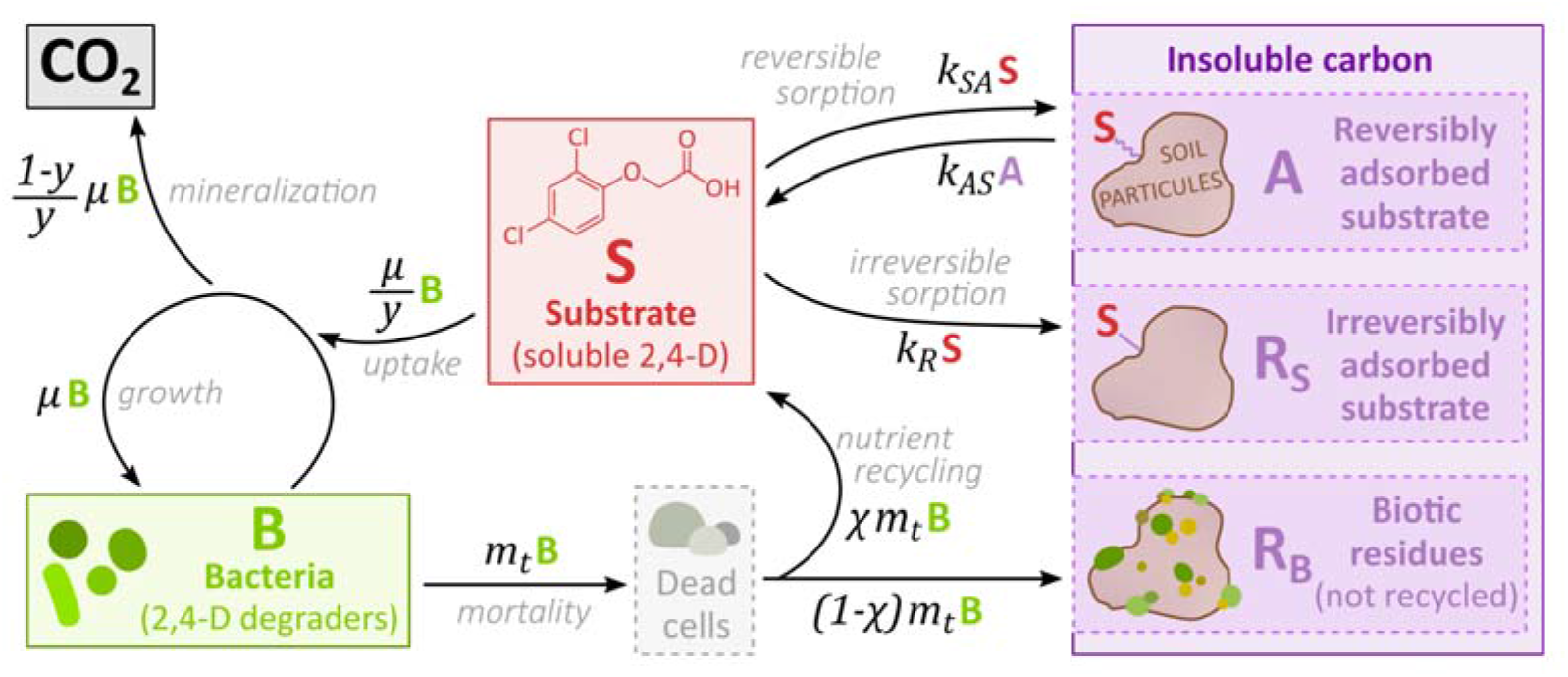
Graphical representation of the biochemical model and carbon fluxes identified by the arrows. Under low substrate concentrations S, the specific uptake rate *(1/y)·µ* becomes equal to *S·(1/y)·µ*_*max*_*/κ*_*M*_, where *(1/y)·µ*_*max*_*/κ*_*M*_ is referred to as the “maximum uptake efficiency”.

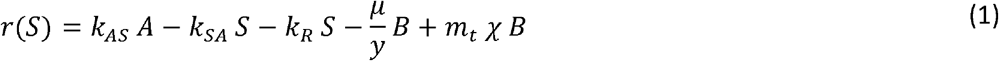

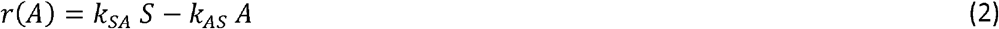

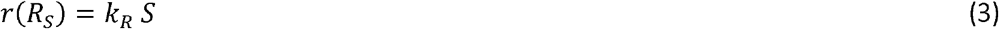

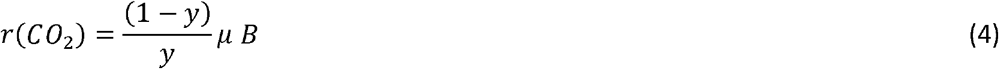

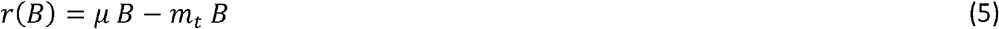

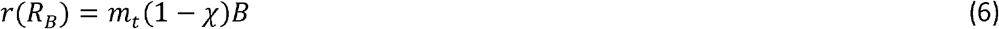

All variable and parameter definitions are listed in **Table 1**. The dynamics of the specific growth rate *µ* are given, for the Monod-based model, by:

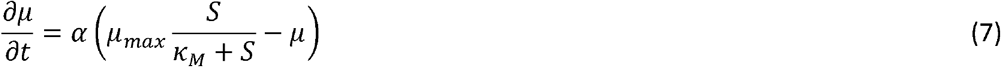

and, for the Contois-based model, by:

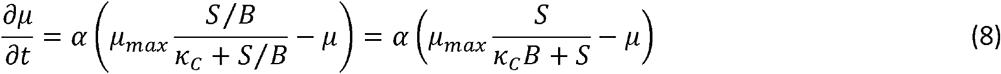

where *µ* = 0 at *t* = 0.

The soluble substrate S is either reversibly adsorbed to soil particles (pool *A*) or irreversibly adsorbed (pool *R*_*S*_) (Eqs. (1), (2), (3)), or taken up by bacteria *B* (Eq. (1)) and metabolized into *CO*_*2*_ (Eq. (4)) and new biomass B (Eq. (5)). *k*_*SA*_ and *k*_*AS*_ are the reversible sorption coefficients. *k*_*R*_ is the irreversible one. Bacteria death occurs at a constant rate *m*_*t*_ (Eq. (5)) and a fraction of the bacterial necromass is considered to return to the soluble substrate pool *S* to account for nutrient recycling (Eq. (1)), while the rest is transformed to biotic residues *R*_*B*_ (Eq. (6)). The nutrient recycling is necessary to adequately predict the late dynamics of mineralization. Its impact on mineralization is only marginal during the first five days. The adsorbed substrate and biotic residues form the pool of insoluble carbon *A + R*_*S*_ *+ R*_*B*_. The substrate *S* is consumed by bacteria *B* according to their specific uptake rate *(1/y)·µ* expressed either by the substrate-dependent Monod growth law (Eq. (7)) (Monod, 1949) or by the ratio-dependent Contois growth law (Eq. (8)) (Contois, 1959). *y* is the yield coefficient and relates the specific uptake rate *(1/y)·µ* to the specific growth rate *µ. µ*_*max*_ is the maximum specific growth rate. *κ*_*M*_ and *κ*_*C*_ are Monod and Contois constants respectively. The effective uptake is delayed by the accommodation rate α, which explicitly takes into account the “memory” effects of the bacteria when adapting to new conditions (Patarinska et al., 2000). This delay is necessary to capture the mineralization lag time at the beginning of the experiments (see **Fig. S6**). Over long time periods (*t* » 1/*α*), *µ* follows the exact expression of the Monod or Contois equations. All modeled pools (*S,B, CO*_*2*_, *A, R*_*S*_ and *R*_*B*_) were expressed as carbon concentrations in µg·g^-1^ (mass of carbon per mass of dry soil) considering a soil water content of 0.205 g·g^-1^ (mass of water per mass of dry soil), a bulk density of the soil column of 1.3 10^3^ g·l^-1^ (mass of dry soil per apparent soil volume) and an average bacterial dry weight of 2.8 10^−13^ g corresponding to 1.49 10^−13^ g of carbon per cell. These values of water content and bulk density were those set up in the experiments, the latter corresponding to a water potential adjusted at -31.6 kPa (pF 2.5). The average bacterial weight was assumed based on Dechesne et al. (2010) and Pinheiro et al. (2015). The water-filled pore space (54%, volume of water per volume of pores) was such that oxygen was not considered a limiting factor for 2,4-D degradation.

### 2.3. Reactive transport model

The transport model is based on the diffusion model of Babey et al. (2017) to which advective-dispersive processes explored in the experiments of Pinheiro et al. (2018) are added. Bacterial leaching out and dispersion were observed only in the percolation experiments while the substrate was also reported to diffuse. Hydrodynamic leaching and dispersion were modeled independently, as they result from, respectively, bypass flow through large pores and complex hydrodynamic dispersion processes coming not only from usual flow mechanisms but also from large saturation variations and local redistribution of moisture in the pore network. Due to the lack of adequate experimental data to characterize the details of the dispersion process, we applied a simple isotropic dispersion coefficient. Complementary numerical simulations show that other anisotropic dispersion parameterization are only weakly sensitive (**Fig. S3**). Bacterial and substrate transports were described with the same advective and dispersive parameters. This assumption did not significantly alter the results (**Fig. S4**). Coupled to the equations of the bioreactive model ((1)-(8)), the full reactive transport model is given by:

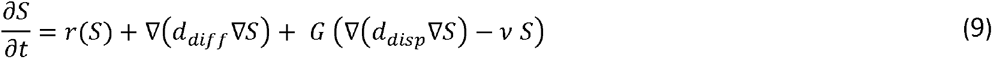

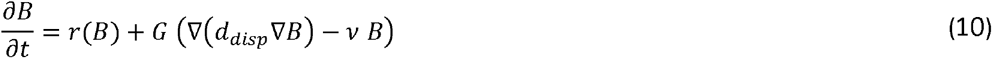

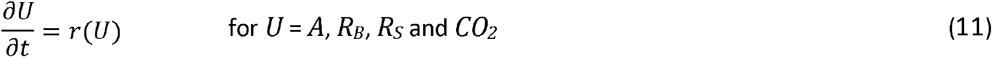

where *d*_*diff*_ is the effective molecular diffusion coefficient of *S, d*_*disp*_ is the effective hydrodynamic dispersion coefficient of *S* and *B* and *ν* is their leaching rate. Note that the dispersion coefficient *d*_*disp*_ mostly affected the spreading of bacteria, given that substrate was mainly spread by diffusion, as noted in section 2.3 and confirmed by consistent results from equivalent models without hydrodynamic dispersion of *S* (**Fig. S5**). Effective diffusion and dispersion processes were assumed to be isotropic and uniform at the column-scale. Dispersion and leaching were active only during the observed 1-hour percolation events at days 0, 3 and 6 as controlled by the function *G* defined as:

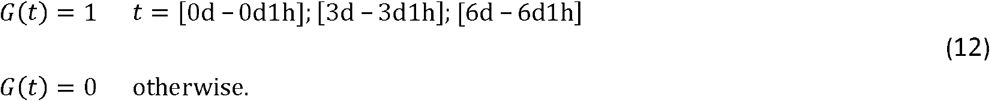

No-flow boundary conditions were imposed at the edges of the soil core (∇*S* = 0 and ∇*B* = 0) during periods outside of the percolation events. The transient evolutions of the water content and their effects on concentrations were not considered because of the short duration of the percolation events (1 h) and the absence of detectable effects on the experimental mineralization curve around the percolation events (**Fig. 1D**). Hydration conditions were considered constant, constrained by the water potential adjusted to -31.6 kPa. No bacterial mobility was observed in the hydrostatic experiments, suggesting that the bacterial mobility observed in the percolation experiments resulted primarily from hydrodynamic dispersion.

Carbon pools concentration dynamics were simulated on a 3 × 6 × 6 regular mesh grid. Although the shape of the grid was slightly different from that of the cylindrical soil-core, it did not have any observable impact (Babey et al., 2017). We recall that substrate and bacteria were initially co-located in the same cube(s). Each cube was considered to be physically, chemically and biologically homogeneous. Diffusion and dispersion were simulated using a finite-difference scheme (Iserles, 2009) and coupled with the bioreactive model, itself solved by the 4^th^ order Runge-Kutta integration method function of MATLAB (Shampine and Reichelt, 1997). The coupling of transport and bioreactive models was achieved with a sequential non-iterative operator-splitting method, in which the equations are resolved within each time step in a sequence of one transport step followed by one bioreactive step (Carrayrou et al., 2004; Lagneau and van der Lee, 2010). The time steps were smaller than the characteristic diffusion and reaction times to avoid any coupling issues.

### 2.4. Exploratory screening

Parameters and their values are listed in **Table 1**. Sorption parameters and the diffusion coefficient were set at values that were calibrated and validated by Babey et al. (2017) in independent experiments without degradation. The mortality rate and the nutrient recycling yield were also kept at the values calibrated in Babey et al. (2017) as they were considered to be well constrained by the residual mineralization dynamics of the homogeneous hydrostatic experiment (**Fig. 1D**). The four biological parameters primarily involved in the biological response of bacteria to the concentration of substrate were determined to be *(1/y)·µ*_*max*_, *α, B(t=0)* and either *(1/y)·µ*_*max*_*/κ*_*M*_ for the Monod-based model or *(1/y)·µmax/(B(t=0)·κC)* for the Contois-based model. Each of these four parameters were sampled over 7 logarithmically-distributed values within the theoretically and physically relevant ranges given by Babey et al. (2017), and all possible combinations of values were screened (**Table S2**). We recall that the “maximum uptake efficiency” *(1/y)·µ*_*max*_*/κ*_*M*_ characterizes the specific bacterial uptake of substrate at the lowest substrate concentration (Button, 1991), while the maximum specific uptake rate *(1/y)·µ*_*max*_ characterizes the bacterial uptake at the highest substrate concentration. Note that the uptake yield y was fixed at the value calibrated by Babey et al. (2017) with a high degree of certainty. The initial maximum uptake efficiency *(1/y)·µ*_*max*_*/(B(t=0)·κ*_*C*_) in the Contois-based model was screened in the same range as *(1/y)·µ*_*max*_*/κ*_*M*_. The accommodation rate α of the degrader response ranged from a negligible delay of few minutes (α = 934 d^-1^) to a prolonged delay of around 10 days (*α* = 9.34 10^−2^ d^-1^). *B(t=0)* values were screened around the initial experimental measurements of the *tfdA* gene copy number, assuming that one *tfdA* sequence corresponded to one bacterium. They ranged over two orders of magnitude to account for the uncertainty of the conversion of *tfdA* copy number into alive 2,4-D degraders (Bælum et al., 2006, 2008). Bacterial density in the uptake efficiency expression will also be expressed in g·l^-1^ (mass of bacteria per volume of water) for a more direct comparison with the relevant literature.

The spatial distribution of bacteria observed at the end of the experiments could not be used to determine the effective dispersion coefficient *d*_*disp*_ (**Fig. S2**). While they qualitatively ascertained that bacteria spread orthogonally to the percolation direction, experimental data were not sufficiently resolved to be used quantitatively. The dispersion coefficient was thus screened over 10 values ranging from no dispersion (*d*_*disp*_ = 0) to complete instant homogenization of the soil core (*d*_*disp*_ = inf) (**Table S2**). The effective diffusion coefficient *d*_*diff*_ had been calibrated independently from percolation conditions (Pinheiro et al., 2015; Babey et al., 2017). The leaching rates ν were determined based on the experimental masses of leached ^14^C (Pinheiro et al., 2018) (**Table 1**). Detailed values for the screened parameters are listed in Table S2.

### 2.5. Model to data comparison

The comparison between the results of the model and the experimental data was based on the core-scale data of mineralization deduced from the carbon mass m_CO2_ of ^14^CO_2_ emissions:

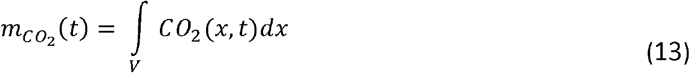

with *V* the volume of the soil cores. Mineralization at a given time t was expressed as the carbon mass of cumulated ^14^CO_2_ emissions 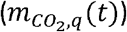. per initial carbon mass of ^14^C-substrate *S* (*m*_*S,q*_(*t* =0) where the index *q* identifies the experiment at hand. Indices *1, 2, 3* and *4* are respectively given to the homogeneous hydrostatic, heterogeneous hydrostatic, homogeneous percolation and heterogeneous percolation experiments. Data-to-model adequacy was assessed for each of the experiments by a classical root-mean-square evaluation function *J*_*q*_ comparing the modeled mineralization of Eq. (4) to the measured mineralization at the *n*_*q*_ available sampling times *t*_*i*_:

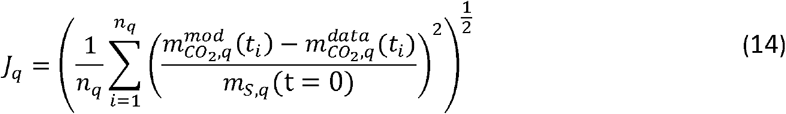

Discrepancies over the full set of experiments J_1234_ were thus expressed as:

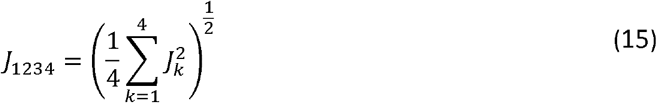

Following the systematic parameter screening described in section 2.5, the parameter set minimizing *J*_*1234*_ was determined and referred to as the set calibrated on both hydrostatic and percolation experiments. The measurement errors were in average 1.7 times higher in the percolation experiments than in the hydrostatic experiments. This was assumed to be due to differences in experimental setup between the two sets of experiments of Pinheiro et al. (2015, 2018). This error difference contributed to limit the weight of the percolation experiments when determining the best-fitting parameter set over the whole set of experiments (*J*_*1234*_). We made the choice to give an equal weight to all experiments by only taking into account the average CO_2_ values.

## 3. Results

### 3.1. Model calibration

The calibration of the bioreactive transport model carried out using only the hydrostatic experimental data (Babey et al., 2017) led to a minimal discrepancy between data and model of *J*_*12*_ = 0.023 (**Fig. 3-A1** and **A2**). This pre-existing parameterization was used to provide blind predictions of the percolation experiments, with the effective dispersion coefficient *d*_*disp*_ as an additional fitting parameter. It gave a reasonable prediction of mineralization in the homogeneous percolation experiment (*J*_*3*_ = 0.038, **Fig. 3-A3**) but failed in the heterogeneous percolation experiment (*J*_*4*_ = 0.151, **Fig. 3-A4**), regardless of the dispersion coefficient values. The smallest discrepancy *J*_*4*_ was surprisingly obtained without any bacterial dispersion (*d*_*disp*_ = 0) in contradiction with the bacterial spread observed in the experimental data (**Fig. S2**). The final predicted mineralization was highest when bacteria remained aggregated close to the initial location of the substrate. The highest predicted mineralization was however four times lower than the experimental data. The large gap between the experimental data and the modeled scenario suggests that bacterial proximity to the initial substrate location is not the underlying explanatory mechanism for the high mineralization rates. On the contrary, it suggests that mineralization might rather be increased by the dispersion of bacteria towards more diluted substrate concentrations, and that the identified bacterial traits do not match this increase of mineralization with dispersion.

**Fig. 3.**
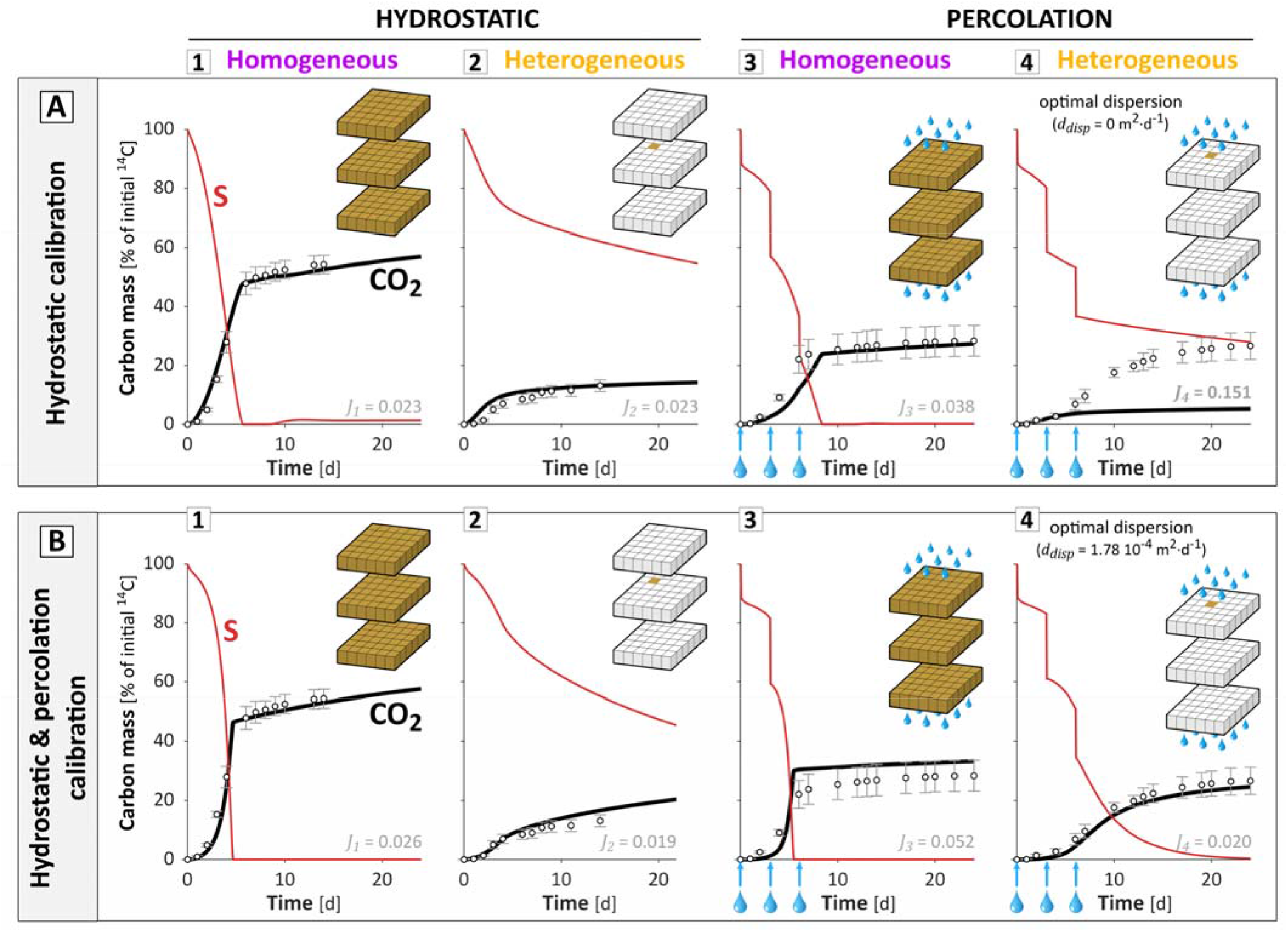
Mineralization dynamics predicted with the Monod-based model calibrated on the hydrostatic experiment only (**A**) and on both hydrostatic and percolation experiments (**B**). The related experimental setups are indicated in the top right corner of each graph. The agreement between experiments and model is indicated by the value of discrepancy *J* displayed on top and can be visually assessed by the proximity between the black line and the dots representing respectively the model results and experimental data. The red line refers to the carbon mass of substrate remaining in the soil core. In the percolation experiments (**A3**,**4** and **B3**,**4**), around 51% of the initial mass of ^14^C was lost through leaching at each percolation events (t = 0, 3 and 6 days, blue arrows). The carbon balance among the different pools is detailed in **Fig. S7**. Note that the reversible sorption eventually accounted for less than 2% of the initial carbon mass and therefore did not significantly alter the results.

In order to investigate the capacity of the reactive transport model to fit both hydrostatic and percolation experimental data, the biological parameters *((1/y)·µ*_*max*_*/κ*_*M*_, *(1/y)·µ*_*max*_, *α, B(t=0))* and the dispersion coefficient (*d*_*disp*_) were calibrated on both hydrostatic and percolation experiments following the screening approach given in section 2.4 to minimize J_1234_. The mineralization dynamics were adequately predicted in all four experiments with the biological parameter set giving the lowest overall discrepancy (*J*_*1234*_ = 0.032) and a non-zero dispersion coefficient (*d*_*disp*_ = 1.78 10^−4^ m^2^·d^-1^) (**Fig. 3, Table 2**). The non-zero dispersion coefficient indicates that the calibrated model accounts for a positive impact of bacterial dispersion on degradation. The model results suggest that this effect is necessary to successfully predict the high degree of degradation in the experimental data. Compared to the parameters calibrated only using the hydrostatic experiments, the parameter set calibrated on both hydrostatic and percolation experiments also displayed a much higher maximum uptake efficiency *(1/y)·µ*_*max*_*/κ*_*M*_ = 26.5 g·µg^-1^·d^-1^ (mass of dry soil per mass of bacterial carbon per unit of time) (Table 2). The systematic exploration of the parameter space showed that high maximum uptake efficiency was a common feature of the 1% best-fitting parameterizations over both hydrostatic and percolation experiments (smallest *J*_*1234*_), with values of 159 and 26.5 g·µg^-1^·d^-1^, corresponding respectively to 1.73 10^4^ and 2.89 10^3^ l·g^-1^·d^-1^ (volume of water per mass of bacteria per unit of time). It underlines the essential role of the maximum uptake efficiency for modulating the impact of dispersion on degradation, further detailed and explained in section 3.2.3.

**Table 2.**
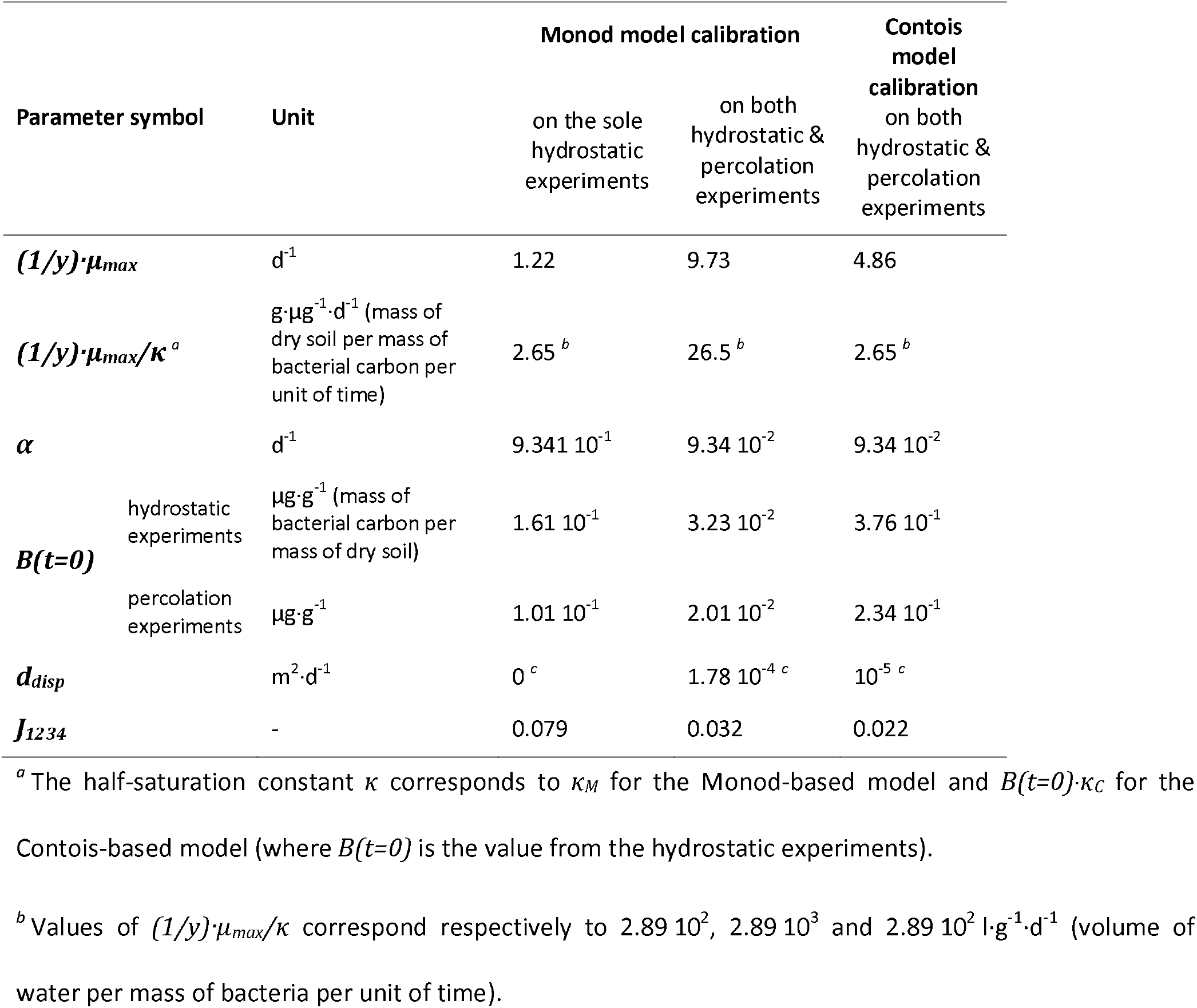
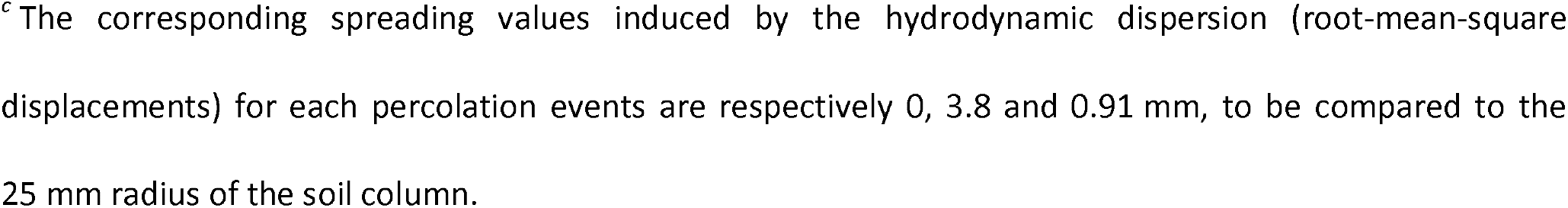
Parameters for the Monod-based model calibrated by the screening approach (section 2.4) on the hydrostatic experiments only (Babey et al., 2017) and on both hydrostatic and percolation experiments, and for the Contois-based model calibrated on both hydrostatic and percolation experiments.

The corresponding spreading values induced by the hydrodynamic dispersion (root-mean-square displacements) for each percolation events are respectively 0, 3.8 and 0.91 mm, to be compared to the 25 mm radius of the soil column.

### 3.2. Analysis of the controls exerted on degradation by substrate dilution and bacterial density

The effect of dispersion on degradation differed greatly between the two calibrated sets of biological parameters described in section 3.1. We therefore conducted a more systematic investigation of the coupled impact of bacterial dispersion and bacterial traits on degradation, revealing its control by substrate dilution and bacterial density.

#### 3.2.1 Impact of dispersion on degradation

We used the mineralization at the end of the experimental time (day 24) as a proxy for degradation and determined its sensitivity to dispersion, as a function of the parameterization of bacterial traits. Fig. 4 shows the impact of the dispersion coefficient *d*_*disp*_ on the final predicted mineralization for the two calibrated biological parameter sets, all other parameters being kept constant (thick red and blue lines). For the biological parameter set calibrated on hydrostatic experiments, the final mineralization decreased monotonically with dispersion (**Fig. 4**, red line). For the parameter set calibrated on both hydrostatic and percolation experiments, the final mineralization first increased, reached a maximum around *d*_*disp*_ ≈ 10^−4^ m^2^·d^-1^ and then decreased (**Fig. 4**, blue line). These two kinds of behaviors were observed regardless of the parameters α, *(1/y)·µ*_*max*_ and *B(t=0)* as long as *(1/y)·µ*_*max*_*/κ*_*M*_ remained the same (Fig. S8). The non-monotonic impact of dispersion on degradation highlights the existence of an optimal bacterial dispersion for which mineralization is the highest. The comparison between the red and blue lines on **Fig. 4** suggests that the optimal dispersion value depends on the bacterial uptake efficiency. Note that, although the optimal dispersion value varied with time due to the spatial dynamics of both bacteria and substrate (**Fig. S9**), it tended towards a limit that was mostly reached within 4 to 7 days and is thus represented at day 24 on **Fig. 4**.

**Fig. 4.**
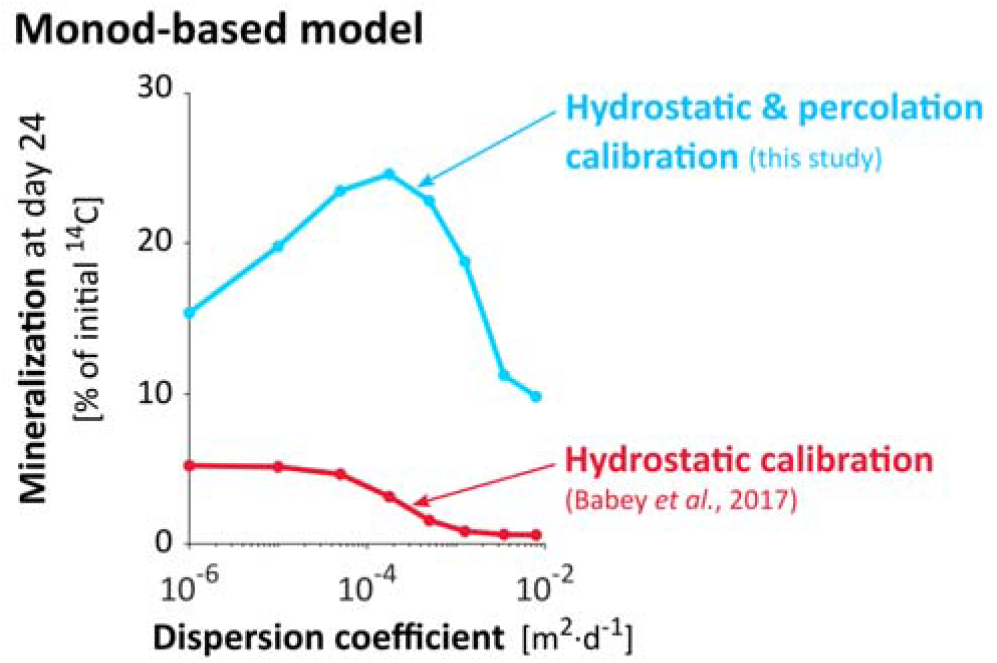
Influence of the dispersion coefficient *d*_*disp*_ on mineralization predicted at day *24 m*_*CO2*_*(t=24)* for the biological parameter set calibrated on the sole hydrostatic experiments (**A**, thick red line) and on both hydrostatic and percolation experiments (**B**, thick blue line). Note that for the model calibrated on both hydrostatic and percolation experiments, the value of *d*_*disp*_ leading to the highest final mineralization (*d*_*disp*_ = 1.78 10^−4^ m^2^·d^-1^, thick blue line) is also equal to its calibrated value leading to the best adequacy with mineralization kinetics (**Table 2**).

#### 3.2.2 Double control of degradation by substrate dilution and bacterial density

The non-monotonic effect of bacterial dispersion on degradation is an unusual and key feature of the model calibrated on both hydrostatic and percolation experiments. In the following we will present an explanation for how such relationships between dispersion and degradation could arise, resulting from a non-monotonic spatial substrate profile, itself derived from the respective effects of substrate dilution and bacterial density. In the model, the instant exposure of bacteria to their substrate is maximal if all the bacteria are located inside the voxel(s) with the highest substrate concentration. In the hydrostatic calibrated parameter set, the profile of substrate concentration primarily resulted from its initial heterogeneity (bell-shape red curve on **Fig. 5A** and pseudo bell-shape red curve on **Fig. 5B**). The flux of substrate reaching each bacterium was therefore mostly determined by the distance between the bacterium and the initial location of substrate. The exposure of a single bacterium to the substrate decreased with its distance from the substrate initial location. This effect is referred to as “substrate dilution”. In these cases (**Fig. 5A** and **B**), mineralization was mainly regulated by substrate dilution, and therefore reduced by bacterial dispersion (**Fig. 4**, blue line). However, for the parameter set calibrated on both hydrostatic and percolation experiments, local degradation by aggregated bacteria reshaped the substrate spatial profile, thus critically changing the voxel(s) with the highest substrate concentration. The bacteria aggregated at their initial location consumed the substrate much faster than it was replenished by backward diffusion and dispersion, creating a critical inversion of the substrate gradient, which led to an intra-population competition for substrate (**Fig. 5C**). The competition was critical for bacterial densities as small as 3.5 10^−3^ g·l^-1^ (**Fig. 5C**). In contrast, the dispersion of bacteria reduced competition by diluting the highest bacterial densities, thus flattening the substrate gradient inversion induced by bacterial local degradation, resulting in a better overall exposure of bacteria to the substrate concentrations, and thus an enhanced mineralization (**Fig. 5D**). In these cases (**Fig. 5C** and **D**), mineralization was mainly regulated by bacterial density. This relation between the bacterial density and the limitation of their exposure to the substrate is not instantaneous and is mediated by the substrate concentration. This is expressed in the model equations through the dependence of bacterial activity *µ(t)* on substrate concentration *S(t)* (Eq. (7)) and the dependence of the substrate concentration *S(t)* on degradation *µ(t)·B(t)* (Eq. (1)), within each voxel. However, when bacterial dispersion was too great, substrate dilution became the dominant control again. This suggests that an optimal bacterial spatial spread exists for which the dilution of substrate is compensated by the dilution of high local bacterial densities. The modeled scenario illustrated by the two calibrated parameter sets were also observed for most of the other parameter sets. The optimal dispersion coefficient for the 300 best-fitting parameterizations to both hydrostatic and percolation experiments (smallest *J*_*1234*_ values) was on average *d*_*disp*_ ≈ 2 10^−5^ m^2^·d^-1^ (**Fig. S10**), corresponding to a root-mean-square displacement of bacteria of 1.5 to 3.5 mm during each percolation event.

**Fig. 5.**
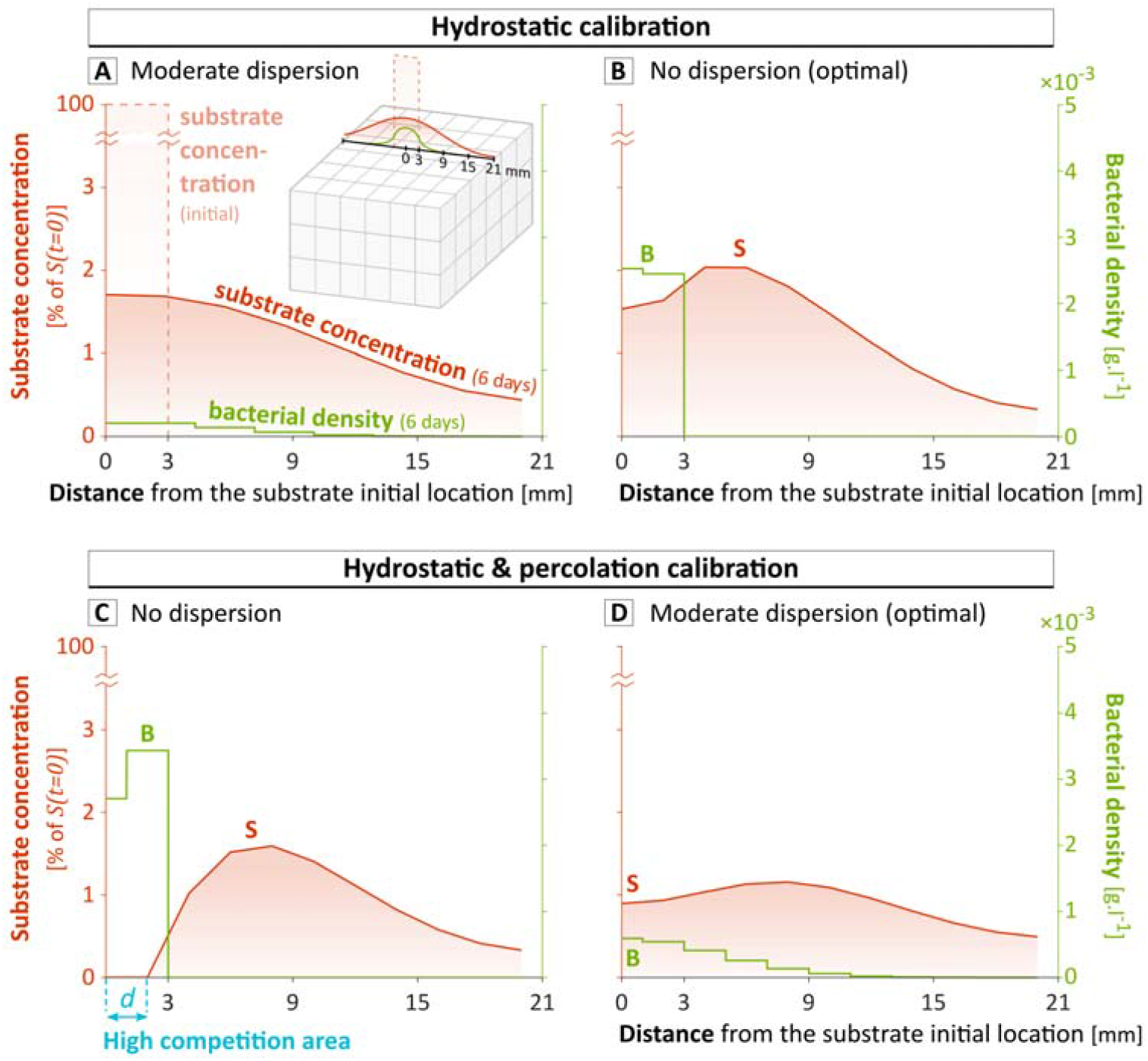
Predicted substrate and bacterial spatial concentration profiles after6 days of diffusion and dispersion in the conditions of heterogeneous percolation experiment, in which bacteria and substrate are initially located exclusively in the central cube (between 0 and 3 mm). Results are simulated on a 9 × 18 × 18 grid obtained by subdividing the 3 × 6 × 6 grid used for the screenings. The results are represented for the parameter set calibrated using only the sole hydrostatic experiment, either with a moderate dispersion (*d*_*disp*_ = 1.78 10^−4^ m^2^·d^-1^) (**A**) or with the calibrated dispersion (no dispersion) (**B**), and for the biological parameter set calibrated on both hydrostatic and percolation experiments, either without dispersion (**C**) or with the calibrated dispersion (*d*_*disp*_ = 1.78 10^−4^ m^2^·d^-1^) (**D**). On one hand, bacteria are exposed to smaller substrate concentrations if they are far from the source (right part of the substrate concentration profiles). On the other hand, bacteria undergo competition if they are too close from each other (left part of the substrate concentration profiles). In (**C**), the bacteria aggregated below d consume the substrate faster than it is replenished by backward diffusion and dispersion. The total number of bacteria within the whole soil column at day 6 is similar in (**A**), (**B**), (**C**) and (**D**), respectively equal to 6.0 10^5^, 9.5 10^5^, 11.5 10^5^ and 11.3 10^5^. The final mineralization at day 24 is however strongly different between scenario, reaching respectively 3.2%, 5.3%, 9.1% and 24.7% of the initial mass of ^14^C.

#### 3.2.3 Effect of bacterial uptake efficiency on the impact of dispersion on degradation

A non-monotonic substrate concentration profile only occurs when bacterial degradation locally depletes the substrate faster than it is replenished by diffusion. This area of high local competition for substrate results from either high local densities of bacteria or high competitiveness or both. Bacterial competitiveness is related to their maximum uptake efficiency *(1/y)·µ*_*max*_*/κ*_*M*_, which also describes their capacity to maintain their activity and growth under dilute substrate concentrations (Healey, 1980; Button, 1991; Lobry et al., 1992). Bacteria with high maximum uptake efficiency are thus expected to benefit more from dispersion. **Fig. 6** shows the optimal dispersion coefficient as a function of the maximum uptake efficiency, with all other parameters equal to those of the model calibrated on both hydrostatic and percolation experiments. The optimal dispersion coefficient, defined as the dispersion coefficient maximizing the final mineralization, increased with the maximum uptake efficiency. For small maximum uptake efficiencies of 30 l·g^-1^·d^-1^ and below, mineralization was highest in the absence of dispersion, suggesting a regulation dominated by substrate dilution. For larger maximum uptake efficiencies, dispersion impacted positively mineralization, suggesting that degradation shifted from being regulated by substrate dilution to being regulated by bacterial densities, as bacteria were both more prone to competition between themselves and more efficient under diluted substrate conditions. In other words, the proximity to other bacteria constrained activity more than the proximity to the substrate initial location enhanced it. This combined effect of the maximum uptake efficiency and the bacterial dispersion on degradation was a general relationship common to all parameterizations (**Fig. S11**).

**Fig. 6.**
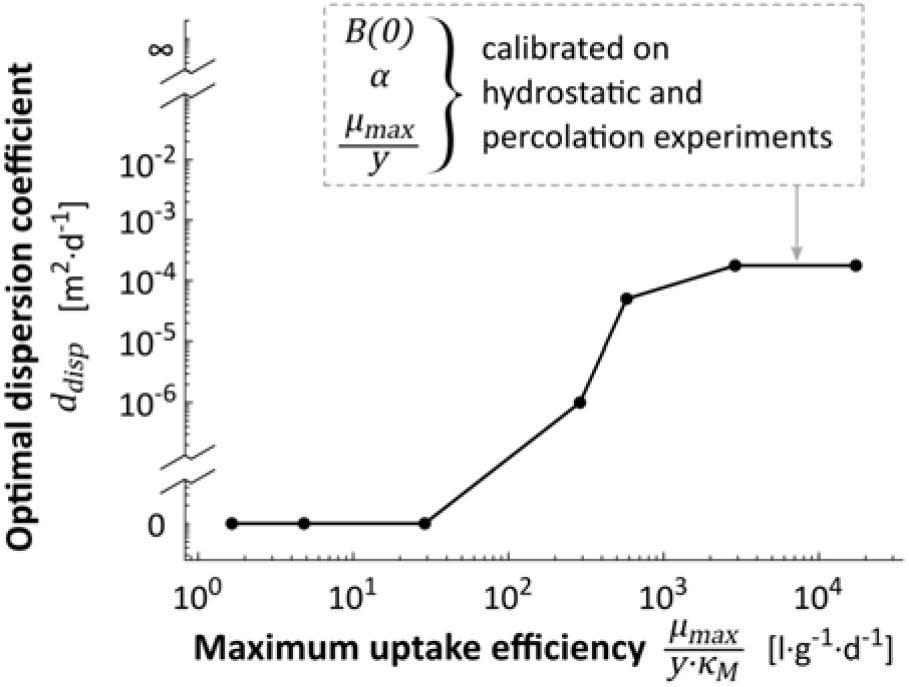
Dispersion coefficient giving the highest predicted mineralization at day 24 as a function of maximum uptake efficiency, all other parameters equal to those of the model calibrated on both hydrostatic and percolation experiments.

### 3.3. The Contois-based model as an alternative to Monod

Given that degradation is regulated by both substrate dilution and bacterial density, and that their relative importance is modulated by bacterial uptake efficiency at the lowest substrate concentration, *(1/y)·µ*_*max*_*/κ*_*M*_, we investigated the relevance of the Contois model by applying the calibration methodology of section 2.4, as used in section 3.1. The interest in the Contois growth law (Eq. (8)) stems from the inclusion of a regulation by density in the expression of the uptake efficiency at the lowest substrate concentration, becoming *(1/y)·µ*_*max*_*/(B(t)·κ*_*C*_).

In comparison with the Monod-based model, the predictions of the experimental observations of Pinheiro et al. (2015, 2019) were facilitated with the Contois-based model, on three levels. First, the Contois-based model captured the degradation dynamics better than the Monod-based model, especially for the 1% best-fitting parameterizations (smallest *J*_*1234*_ values) (**Fig. S12**). The calibrated Contois-based model had an overall discrepancy of *J*_*1234*_ = 0.022 (**Fig. 7**), which was smaller than the lowest value of J_1234_ = 0.032 obtained for the calibrated Monod-based model (**Fig. 3**).

**Fig. 7.**
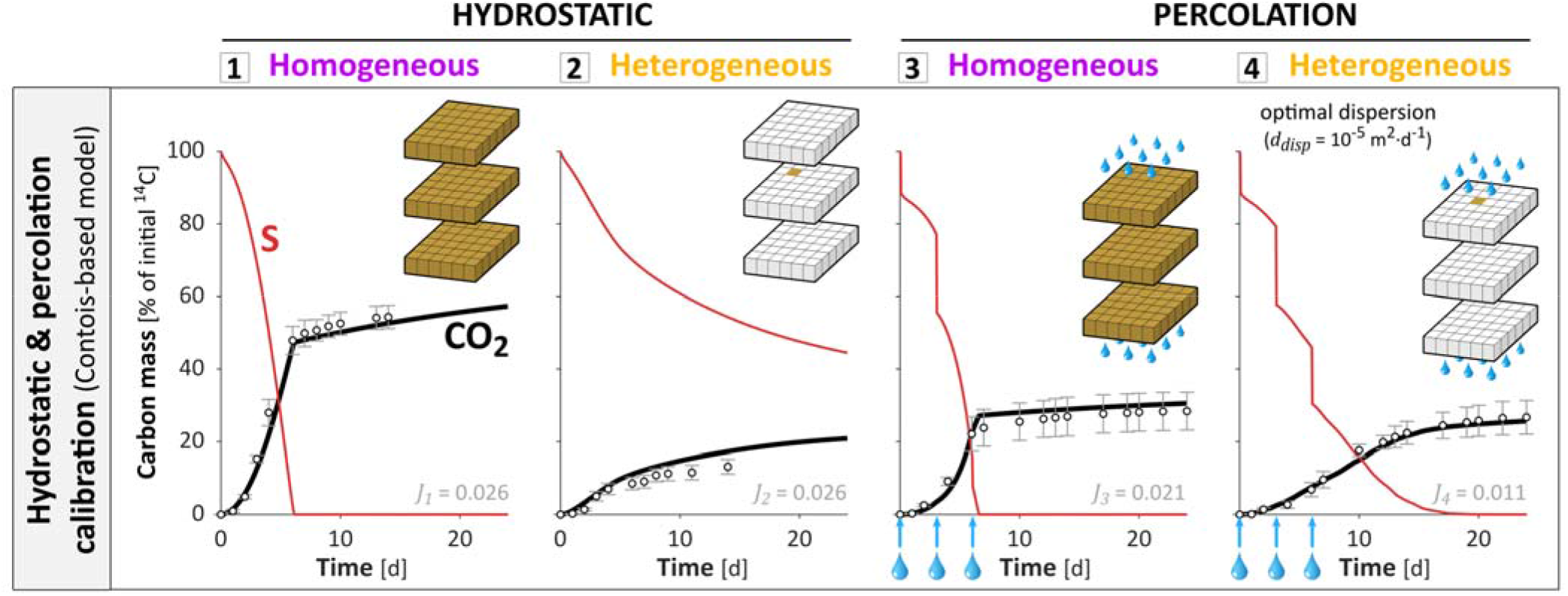
Mineralization dynamics predicted with the Contois-based model calibrated on both hydrostatic and percolation experiments. For representation and legend, see **Fig. 3**. The carbon balance among the different pools is detailed in **Fig. S7**.

Second, the parameter sets that fitted homogeneous experiments also performed well under heterogeneous conditions, as long as the dispersion coefficient *d*_*disp*_ was calibrated as well (**Fig. S13**). It is an important advantage as it confers a better capacity to predict degradation kinetics for heterogeneous and varying distributions, once the model is calibrated in homogeneous conditions, which are more appropriate for the experimental measurement of bacterial parameters. Besides, using a dispersion coefficient value different from the calibrated one weakened the predictions of the mineralization dynamics but not the predictions of the mineralization after 24 days, which remained satisfying regardless of the dispersion coefficient. More precisely, the prediction of the final mineralization became mostly independent of the dispersion coefficient, as shown for the calibrated model (**Fig. 8**). This is because, in the Contois model at low substrate concentrations, the number of active bacteria in a soil volume is exactly counterbalanced by the regulation of their uptake efficiency by population density (Eq. (8)), resulting in limited effects of bacterial spreading on overall mineralization (**Fig. 8**, constant part of the curves).

**Fig. 8.**
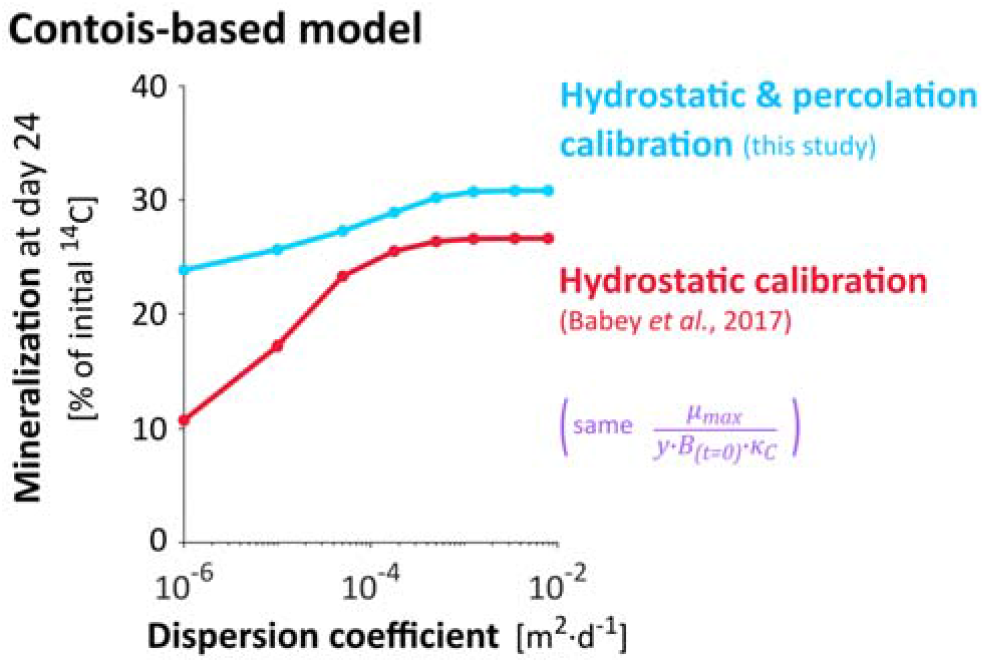
Influence of the dispersion coefficient on mineralization at day 24 for the Contois-based models calibrated on the sole hydrostatic experiments (thick red line) and on both hydrostatic and percolation experiments (thick blue line). For representation and legend, see **Fig. 4**.

## 4. Discussion

### 4.1. Relevance of density control for 2,4-D degradation and soil carbon cycling

#### 4.1.1 Density control of soil oligotroph bacteria

Bulk soil and highly-diluted environments are usually found to be dominated by bacteria with high maximum uptake efficiency, also called oligotrophs (Fierer et al., 2007; Nunan et al., 2020). Their high maximum uptake efficiency differentiates their life-history strategies and conditions their ability to thrive in resource poor environments (Button, 1993), also assimilated to K-strategy (Tecon and Or, 2017), by opposition to copiotrophic bacteria adapted to rich environments (r-strategy). The maximum uptake efficiency values of the 1% best-fitting parameter sets were of the order of 10^3^-10^4^ l·g^-1^·d^-1^ (volume of water per mass of bacteria per unit of time), within the range proposed by Button (1991) to define oligotrophs. Similar or higher maximum uptake efficiency values of the order of 10^4^-10^5^ l·g^-1^·d^-1^ have been reported for soil oligotrophs (Ohta and Taniguchi, 1988; Zelenev et al., 2005). Values up to 1.64 10^5^ have been reported by Tuxen et al. (2002) for 2,4-D degraders in an aerobic aquifer and even greater values might also be possible (see section S5). The high maximum uptake efficiencies predicted in section 3.1 for the best-fitting parameterizations are therefore a plausible bacterial trait among 2,4-D degraders as well as bulk soil bacteria in general. It suggests that density control might be relevant for a component of soil bacteria, which would benefit from dispersion as suggested by **Fig. 6**. The calibrated model has shown in section 3.2.2 that the values of densities from which competition became critical were around 3.5 10^−3^ g·l^-1^, corresponding to 7.5 10^−7^ g·g (mass of bacteria per mass of dry soil), ranging in the low end of usual total soil bacterial densities (Raynaud and Nunan, 2014; Kuzyakov and Blagodatskaya, 2015). This suggests that competition might play a significant role even under the low bacterial densities observed in bulk soils. Reciprocally, the model suggests that competition for substrate between copiotrophic bacteria only appears at much larger population densities, such as those found in soil biofilms (Holden et al., 1997, Or et al., 2007). Interestingly, copiotrophic bacteria have been reported to cohabit with oligotrophic bacteria even in diluted environments (Gözdereliler et al., 2012). Results from the screening suggest that, for densities of copiotrophs as low as for oligotrophs, their impact on overall decomposition in dilution-dominated environments would be much lower due to their poorly adapted uptake efficiency (**Fig. 4A**). Conversely, this striking density regulation might be one of the main limitations of the overall population densities in soils. Note that this density regulation occurs within a single population with homogeneous biological constants. Spatial heterogeneities and low substrate concentrations, common in bulk soil, may indeed shift competition from the inter-population level to the intra-population level (Pfeiffer et al., 2001; Roller and Schmidt, 2015).

#### 4.1.2 A new perspective on Regulatory Gate hypothesis

Density regulation might partially contribute to explain the common paradox of the apparent uncoupling between the overall mineralization of a soil volume and the size of its microbial population (Kemmitt et al., 2008). The rate of soil carbon mineralization remains the same even if 90% of the microbial decomposers are killed. This observation is commonly explained by the Regulatory Gate hypothesis, where mineralization is assumed to be controlled by an abiotic process, such as desorption or diffusion, that limits the availability of the substrate, resulting in mineralization rates that are independent of the degrader abundance. We propose that the density regulation of decomposition in oligotrophic environments may contribute to this phenomenon, through competition for substrate or other biological interactions. In the case of competition-related density regulation, it reduces the dependence of the overall carbon mineralization on degrader abundance, as any increase of population density counterbalances the effect of the increased population size. Note that the involved abiotic process, namely the substrate diffusion backward to bacteria (see section 3.2), is well limiting but only in situations of high bacterial competition.

### 4.2. Relevance of the ratio-dependent Contois model in soils

As argued in section 3.3, ratio-dependence might facilitate decomposition modeling in the soil conditions typical of the experiments analyzed here. The Contois model’s *(1/y)·µ*_*max*_*/κ*_*C*_*B* calibrated in homogeneous conditions might be used in heterogeneous conditions more reliably than the Monod model’s *(1/y)·µ*_*max*_*/κ*_*M*_, at least for soil systems in which the competition for the substrate plays a substantial role within the degrader population. The similarity between *κ*_*M*_ and *κ*_*C*_*B* suggests the need to consider population density when measuring the apparent maximum uptake efficiency of soil bacteria to avoid underestimating it by unintentionally including density regulation. Moreover, the better predictions obtained with the Contois model in the soil conditions represented by the experiments suggest that the Contois ratio-dependence includes not only the effect of competition for substrate at the scale of measurement, but it can also reasonably reflect other density processes such as the spatial variability of bacterial distributions at finer scales related to their high degree of local aggregation in microcolonies (Raynaud and Nunan, 2014). Moreover, ratio-dependence may also include the cumulative effects of ecological interactions other than competition (Sibly and Hone, 2002). Note that the methodological approach used in this study for both Monod and Contois models is based on an effective representation of concentrations and parameters at the mm- to cm-scale of measurements. These effective concentrations and parameters conceptually integrate the smaller-scale processes highlighted by other studies (Ebrahimi and Or, 2014; Portell et al., 2018; Tecon et al., 2018). Such microscale processes should be addressed for further generalization beyond the conditions of the soil experiments analyzed here. Despite its advantages, Contois models have also a drawback with the fact that the modeled uptake efficiency of bacteria approaches infinity for low densities, which does not correspond to any physical nor biochemical process (Gleeson, 1994; Abrams, 2015). However, this side effect mostly affects a negligible fraction of the bacteria and the substrate, as it was the case in the soil conditions represented by the experiments.

Further work is required to confront the relevance of the Contois model to other soil systems. To the best of our knowledge, ratio-dependent growth models such as the Contois model have not yet been considered for the modeling of microbial degradation in soils. However, the Contois growth equation is generally accepted to be more appropriate than the Monod equation for modeling immobilized, heterogeneously distributed or mixed microbial cultures (Arditi and Saiah, 1992; Harmand and Godon, 2007), all of which are characteristics of soils. The regulation of individual activity by population density has frequently been justified as a “crowding effect” associated with high population densities leading to competition for substrate (Lobry and Harmand, 2006; Harmand and Godon, 2007; Krichen et al., 2018). However, little is known about possible density regulation when apparent microbial densities are low, as is observed in bulk soil (Raynaud and Nunan, 2014; Kuzyakov and Blagodatskaya, 2015), although some studies have mentioned ratio-dependence in highly-diluted environments such as aquifers (Hansen et al., 2017). As discussed in section 4.1.1, the high maximum uptake efficiencies commonly observed for soil bacteria adapted to oligotrophic environments are relevant to draw attention on the potential significance of density control at low densities in oligotrophic soils, and thus ratio-dependent models, among which the Contois model is a consistent choice.

### 4.3. Hypothetical relationship between bacterial traits and their spatial strategies

Density regulation might be at the origin of a relationship between bacterial oligotrophy, their location in soil and their mobility strategy. Soil copiotroph bacteria have a maximum uptake efficiency mostly between 100 l·g^-1^·d^-1^ (Button, 1991) and 800 l·g^-1^·d^-1^ (Daugherty and Karel, 1994; Zelenev et al., 2005). For copiotrophs with maximum uptake efficiency values below 288 l·g^-1^·d^-1^, bacterial dispersion was largely detrimental to their activity (**Fig. 4** blue line, **Fig. 6**), in agreement with the results of Pagel et al. (2020), suggesting that copiotrophs have more aggregated distributions than oligotrophs. The negligible mineralization even without dispersion **(Fig. 3-A4, Fig. S8**) also highlights the fact that copiotrophs are particularly inefficient at degrading substrates that diffuse in the environment, as also evidenced by Babey et al. (2017). To maintain significant activity, soil copiotrophs are likely to remain immobile in the close surroundings of the substrate source or any immobile substrate, likely attached to surfaces or embedded in EPS matrices. If not, they would be dispersed towards more diluted area where their low maximum uptake efficiency would result in negligible uptake. On the contrary, to survive and develop, soil oligotrophs should be able to easily disperse and escape high competition areas. Given that soil is a poor and heterogeneous environment, this dispersion would be essentially passive (Nunan et al., 2020), through advective processes for example. We therefore suggest the existence of a theoretical relationship between proximity to substrate sources (respectively remoteness), copiotrophy (respectively oligotrophy) and attachment (respectively mobility).

## 5. Conclusions

Heterogeneous distributions of degraders and substrate in soils strongly control soil organic matter degradation through their interactions with the bacterial activity. Taking 2,4-D as a model organic solute substrate for soil bacteria, we investigated the coupled effects of bacteria and substrate distributions on one side and bacterial traits on the other side on substrate degradation. The analysis of published experiments with contrasted spreading conditions of both bacteria and substrate reveals that, in addition to the distance of bacteria from high substrate concentrations, mineralization is also surprisingly limited by the bacterial density even under the low bacterial densities commonly observed in bulk soils. Moreover, the impact of bacterial dispersion on solute substrate degradation can shift from negative to positive depending on the bacterial maximum uptake efficiency. The activity of soil oligotrophs may be mostly regulated by bacterial density rather than by substrate dilution, echoing the population size paradox regularly observed. It follows that the ratio-dependent Contois model might be more relevant to model bulk soil mineralization in the heterogeneous conditions investigated than the substrate-dependent Monod model. To predict the impact of spatial distributions on degradation in oligotrophic soil, and more particularly the impact of bacterial dispersion, we suggest that bacterial densities might be a more useful measurement than the volumes of soil devoid or occupied with bacteria. With respect to the current lack of direct microscale data on microbial processes and distributions, we propose some key perspectives on the bacterial kinetics and distributions.

## Acknowledgements

This work was supported by the Agence Nationale de la Recherche through the project “Soilµ-3D” [grant number ANR-15-CE01-0006] and was also partially supported by the SLAC Floodplain Hydro-Biogeochemistry Science Focus Area (SFA), which is funded by the U.S. Department of Energy (DOE) office of Biological and Environmental Research (BER), Climate and Environmental Sciences Division, under DOE contract No. DE-AC02-76SF00515 to SLAC. The authors thank Jérôme Harmand, Théodore Bouchez, Xavier Raynaud, Tanguy Le Borgne, Claire Chenu and Holger Pagel for insightful discussions.

## Appendix A. Supplementary data

