## Supplementary_Data for "Competition within low-density bacterial populations as an unexpected factor regulating carbon decomposition in bulk soil"

The following supporting information is available for this article:

***S.1. Experimental conditions of soil-column laboratory experiments ..............................................2***

- **Fig. S1**. Set up of the soil column experiments
- **Table S1**. Experimental conditions for the four experiments
- **Fig. S2**. Final experimental millimeter spatial distributions of bacteria

***S.2. Screening ........................................................................................................................5***

- **Table S2**. Detailed parameter values used in the screenings

***S.3. Transport model ...............................................................................................................6***

- **Fig. S3**. Impact of dispersion anisotropy
- **Fig. S4**. Impact of dispersion on mineralization predicted at day 24, depending on the scenario of bacteria leaching
- **Fig. S5**. Impact of dispersion on mineralization predicted at day 24, with or without hydrodynamic dispersion of substrate

***S.4. Supplementary results ...................................................................................................10***

- **Fig. S6**. Main modeling results without the explicit bacterial accommodation rate $\alpha$
- **Fig. S7**. Carbon mass balance for the Monod based and the Contois based models
- **Fig. S8**. Influence of the dispersion coefficient on mineralization predicted at day 24 for the different calibrated biological parameter sets
- **Fig. S9**. Convergence of optimal bacterial spreading with time
- **Fig. S10**. Relevance of bacterial spreading
- **Fig. S11**. Impact of maximum uptake efficiency on the optimal dispersion coefficient
- **Fig. S12**. Adequacy between simulations and experiments under Monod-based and Contois-based models
- **Fig. S13**. Predictive capacities of Monod-based and Contois-based models for aggregated and dispersive conditions

***S.5. Theoretical collision frequency between bacteria and solute substrate ..................................22***

### *S.1. Experimental conditions of soil-column laboratory experiments*

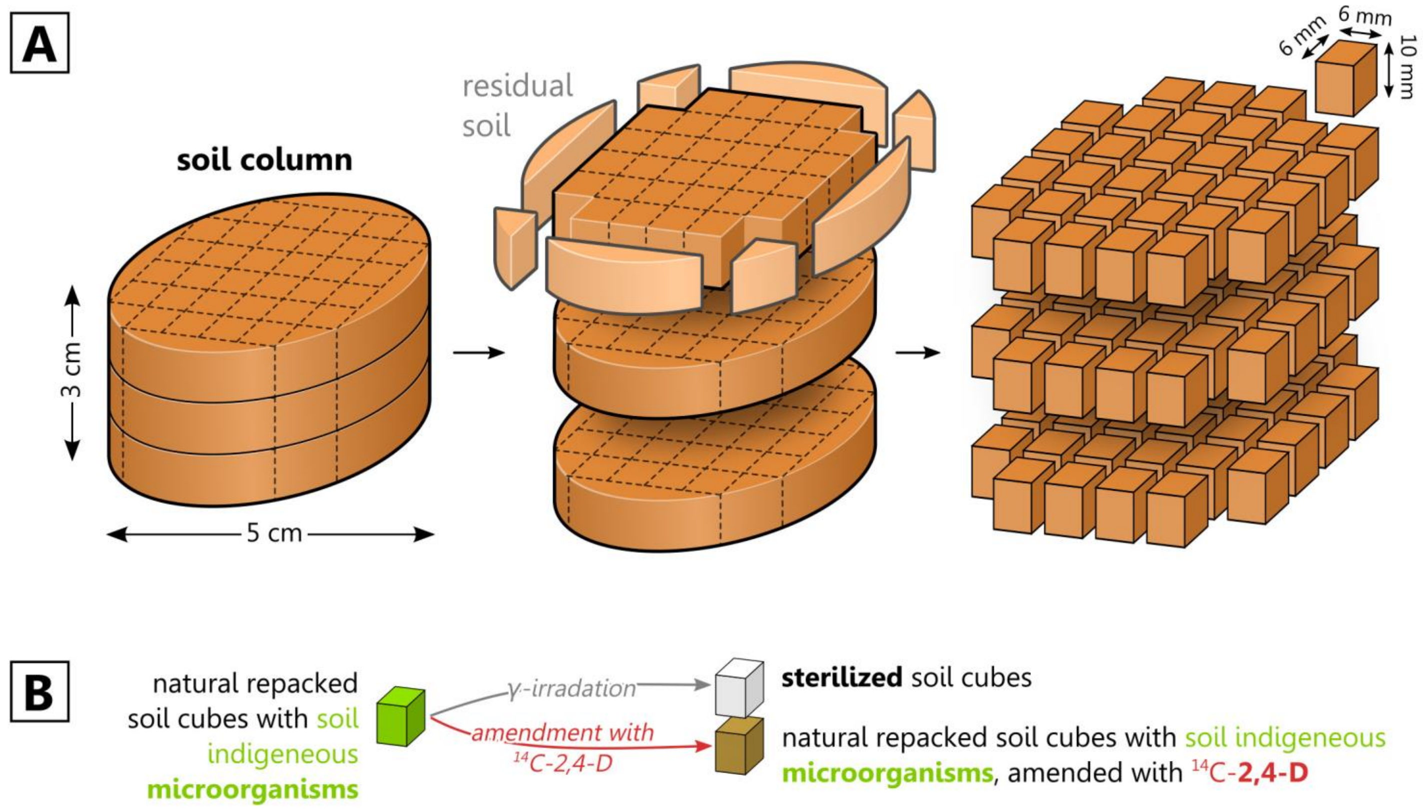


**Fig. S1**. Set up of the soil column experiments. (**A**) Decomposition of cylindrical soil columns into 96 soil cubes. (**B**) Schematic representation of the different types of soil cubes. Soil columns were 5 cm large and 3 cm high and were composed of 0.6 g soil cubes of approximatively 6 mm × 6 mm × 10 mm. Each cube was packed with 2-3.15 mm soil aggregates sieved from a soil sampled from the ploughed layer (0-30 cm) of an agricultural field cultivated without any 2,4-D application for the last 15 years. The different types of soil aggregates were obtained by irradiation and amendment prior to being repacked into the soil column. The water potential of the soil column was kept at -31.6 kPa (pF 2.5), corresponding to a water content of 0.205 $g{\cdot}g^{-1}$ (mass of water per mass of dry soil) and a pore saturation of 54% (volume of water per volume of pores). The apparent density of the soil column was 1.3 $10^3$ $g{\cdot}l^{-1}$ (mass of dry soil per apparent soil volume).

**Table S1**.

Experimental conditions for each experiment. Diffusion applies only to the substrate ($^{14}C$-2,4-D), whereas hydrodynamic dispersion applies to both substrate and bacterial degraders. Diffusion and dispersion reshape the column-scale distributions of mobile species only in heterogeneous experiments, as indicated by the grey shading.

| | | Effective diffusion coefficient $d_{diff}$ [$m^2 \cdot d^{-1}$] | Initial degraders density $B(t=0)$ [$g^{-1}$] (number of tfdA gene copies per mass of dry soil) | Initial $^{14}C$-2,4-D concentration $S(t=0)$ [$g \cdot g^{-1}$] (mass of 2,4-D per mass of dry soil) | Percolation events |
|---|---|---|---|---|---|
| **Hydrostatic experiments** | **homogeneous** | 0.6 $10^{-5}$ [a] | 4.65 $10^{5}$ [b] | 1.9 $10^{-6}$ [c] | no percolation |
| | **heterogeneous** | 0.6 $10^{-5}$ [a] | 4.65 $10^{5}$ [b] | 1.9 $10^{-6}$ [c] | no percolation |
| **Percolation experiments** | **homogeneous** | 0.6 $10^{-5}$ [a] | 2.90 $10^{5}$ [b] | 15 $10^{-6}$ [c] | leaching and hydrodynamic dispersion |
| | **heterogeneous** | 0.6 $10^{-5}$ [a] | 2.90 $10^{5}$ [b] | 15 $10^{-6}$ [c] | leaching and hydrodynamic dispersion |

[a] The values of $d_{diff}$ correspond to the value of 2,4-D effective diffusion coefficient from Babey et al. (2017) calibrated on a 6 × 12 × 12 grid in similar conditions. Note that the diffusion coefficient was calibrated at 1 $10^{-5}$ $m^2 \cdot d^{-1}$ on a 3 × 6 × 6 grid.
[b] $B(t=0)$ values for the hydrostatic and percolation experiments correspond respectively to 2.27 $10^9$ and 1.42 $10^9$ $l^{-1}$ (number of *tfdA* gene copies per volume of water).
[c] $S(t=0)$ values for the hydrostatic and percolation experiments correspond respectively to solute concentrations of 41.9 and 331 $\mu mol \cdot l^{-1}$ (amount of 2,4 D per volume of water)

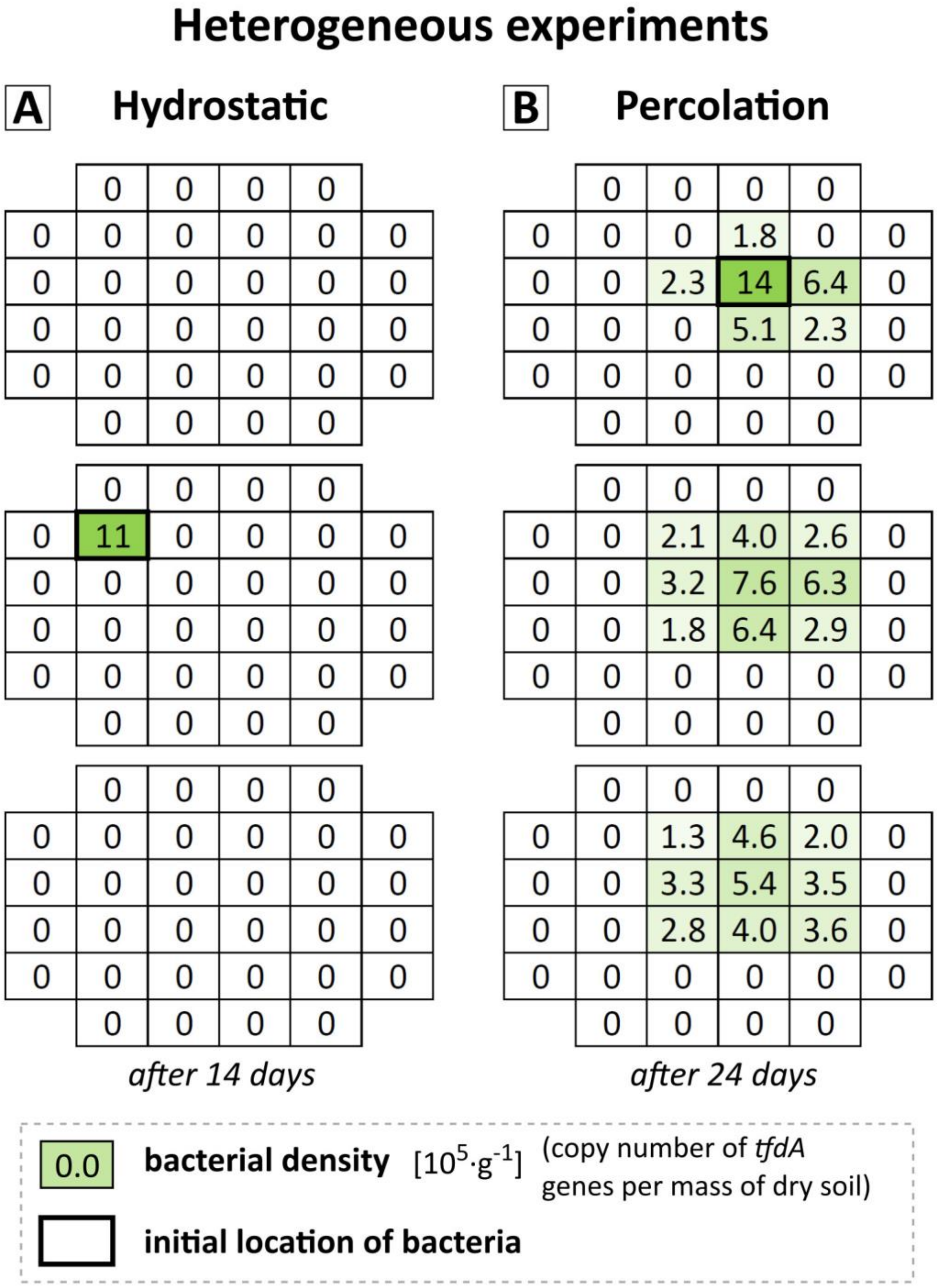


**Fig. S2**. Final experimental millimeter spatial distributions of bacteria, in the heterogeneous experiments (left and right). The 96 experimental cubes are represented on 3 grids corresponding to the 3 layers (top, middle and bottom). The values refer to the final *tfdA* measurements, also assessed by the green color scale. The cell with bold outlines refers to the initial locations of bacteria. The detection limit was 0.12 $10^{5}$ $g^{-1}$ (copies of *tfdA* sequences per mass of dry soil). Adapted from Pinheiro et al. (2018). Permission for reproduction granted by Elsevier.

### *S.2. Screening*

**Table S2**.
Detailed parameter values used in the screenings. Blue cells refer to the parameterization of the biochemical model (Eqs. (1)-(8)) calibrated on the sole hydrostatic experiments by Babey et al. (2017), and numbers in blue refer to the ratio to these values. Green cells refer to the number of copies of *tfdA* gene measured by Pinheiro et al. (2018, 2015). The screened values of the four biological parameters were chosen to be more refined around the values previously calibrated on the sole hydrostatic experiments (blue cells). The values of $d_{disp}$ were chosen to be regularly spaced on a log scale, supplemented by a value for no dispersion ($d_{disp}$ = 0) and a value for instant homogeneous distribution ($d_{disp}$ = inf). The broad range of $\alpha$ (five orders of magnitude) accounted for conditions of negligible delay of few minutes ($\alpha$ = 934 $d^{-1}$) to prolonged delay of around 10 days ($\alpha$ = 9.34 $10^{-2}$ $d^{-1}$). The 2 orders of magnitude range of $B(t{=}0)$ accounted for the uncertainty on the conversion of *tfdA* copy number into alive bacterial degraders (Bælum et al., 2008, 2006).

| Biological parameters | | Unit | Screened values | | | | | | |
|---|---|---|---|---|---|---|---|---|---|
| *Maximum specific uptake rate* $(1/y)\cdot\mu_{max}$ | | $d^{-1}$ | **0.0190** | **0.608** | **1.22** | **2.43** | **4.86** | **9.73** | **19.5** |
| | | ratio | 1/64 | 1/2 | 1 | 2 | 4 | 8 | 16 |
| *Maximum uptake efficiency* $(1/y)\cdot\mu_{max}/\kappa$ [a] | | $g\cdot\mu g^{-1}\cdot d^{-1}$ (mass of dry soil per mass of bacterial carbon per unit of time) | **0.0152** | **0.0442** | **0.265** | **2.65** | **5.31** | **26.5** | **159** |
| | | $l\cdot g^{-1}\cdot d^{-1}$ (volume of water per mass of bacteria per unit of time) | 1.65 | 4.81 | $2.89\ 10^{1}$ | $2.89\ 10^{2}$ | $5.78\ 10^{2}$ | $2.89\ 10^{3}$ | $1.73\ 10^{4}$ |
| | | ratio | 1/175 | 1/60 | 1/10 | 1 | 2 | 10 | 60 |
| *Accommodation rate* $\alpha$ | | $d^{-1}$ | **0.00934** | **0.0187** | **0.0934** | **0.934** | **9.34** | **93.4** | **934** |
| | | ratio | 1/100 | 1/50 | 1/10 | 1 | 10 | 100 | 1000 |
| *Initial degraders population density* $B(t{=}0)$ [b] | Hydrostatic exp | $\mu g\cdot g^{-1}$ | **0.0161** | **0.0323** | **0.0692** | **0.161** | **0.376** | **0.807** | **1.61** |
| | | $g\cdot l^{-1}$ | $1.48\ 10^{-4}$ | $2.96\ 10^{-4}$ | $6.36\ 10^{-4}$ | $1.48\ 10^{-3}$ | $3.45\ 10^{-3}$ | $7.41\ 10^{-3}$ | $1.48\ 10^{-2}$ |
| | | ratio | 1/10 | 1/5 | 1/2.33 | 1 | 2.33 | 5 | 10 |
| | Percolation exp | $\mu g\cdot g^{-1}$ | **0.0101** | **0.0201** | **0.0432** | **0.101** | **0.234** | **0.503** | **1.01** |
| | | $g\cdot l^{-1}$ | $9.24\ 10^{-5}$ | $1.85\ 10^{-4}$ | $3.97\ 10^{-4}$ | $9.24\ 10^{-4}$ | $2.15\ 10^{-3}$ | $4.62\ 10^{-3}$ | $9.24\ 10^{-3}$ |
| | | ratio | 1/10 | 1/5 | 1/2.33 | 1 | 2.33 | 5 | 10 |

| Physical parameter | Unit | Screened values | | | | | | | | | |
|---|---|---|---|---|---|---|---|---|---|---|---|
| *Dispersion coefficient* $d_{disp}$ [c] | $m^2\cdot d^{-1}$ | **0** | **$10^{-6}$** | **$10^{-5}$** | **$5.01\ 10^{-5}$** | **$1.78\ 10^{-4}$** | **$5.01\ 10^{-4}$** | **$1.26\ 10^{-3}$** | **$3.47\ 10^{-3}$** | **$7.94\ 10^{-3}$** | **∞** |
| | mm at each event | 0 | 0.289 | 0.913 | 2.04 | 3.85 | 6.46 | 10.2 | 17.0 | 25.7 | ∞ |

[a] The half-saturation constant $\kappa$ corresponded to $\kappa_M$ for the Monod-based model and $B(t{=}0)\cdot\kappa_C$ for the Contois-based model (where $B(t{=}0)$ is the value from the hydrostatic experiments).
[b] Values of $B(t{=}0)$ were set individually in each experiment according to the initial *tfdA* gene abundance differences measured in experiments, as the number of *tfdA* copies per volume of water was 1.6 times smaller in the percolation experiments (0.0432 $\mu g\cdot g^{-1}$) than in the hydrostatic experiments (0.0692 $10^9$ $\mu g\cdot g^{-1}$).
[c] The effective dispersion coefficient $d_{disp}$ applied only to heterogeneous percolation experiments. Length values below in mm corresponds to the equivalent root mean squared spreading of bacteria and substrate by each irrigation event.

### *S.3. Transport model*

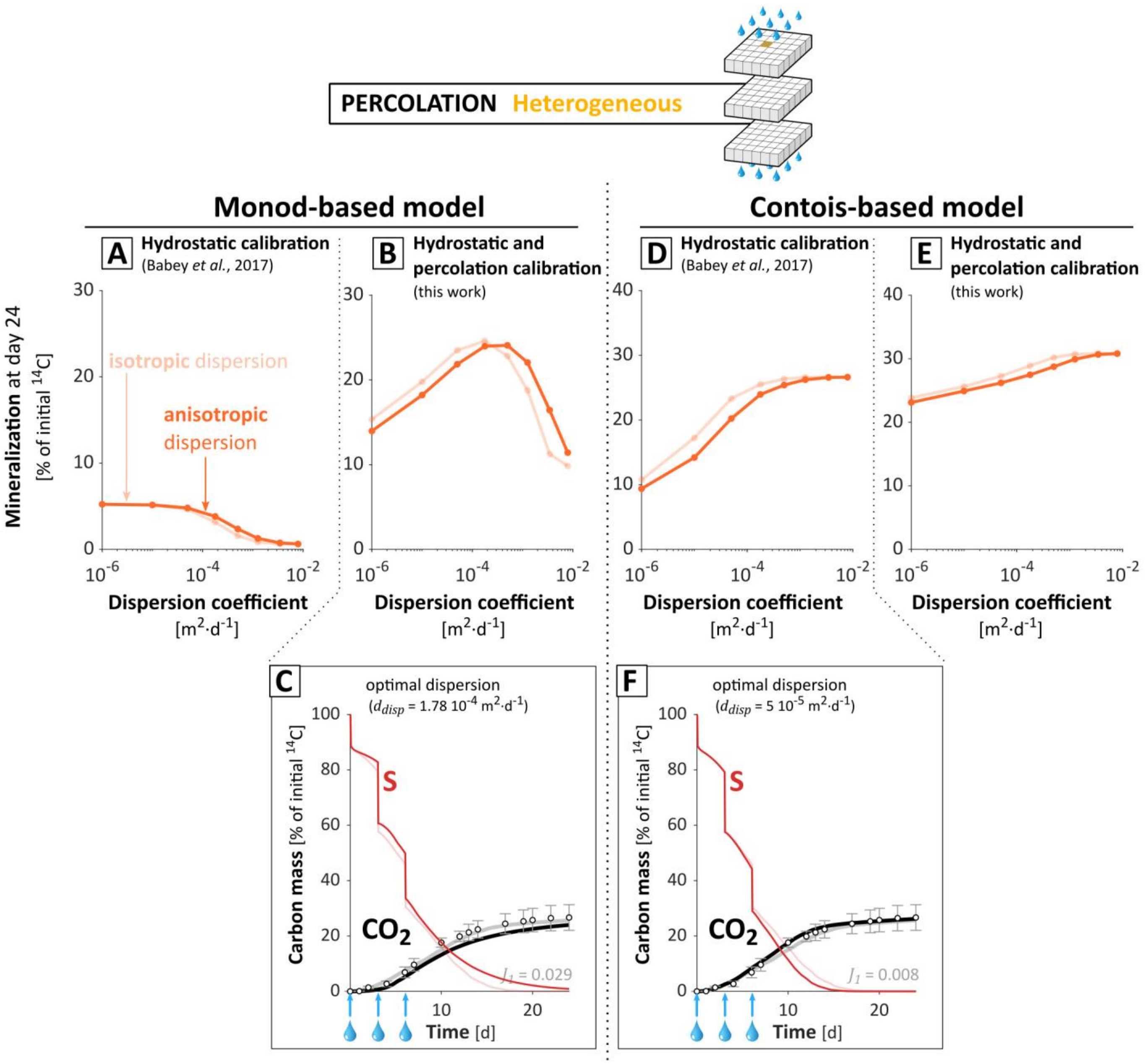


**Fig. S3.** Impact of dispersion anisotropy. (**A**-**B**-**C**) refer to the Monod model and (**D**-**E**-**F**) refer to the Contois model. Figures at the top (**A**, **B**, **D**, **E**) represent the mineralization predicted at day 24 as a function of the dispersion coefficient $d_{disp}$, depending on the anisotropy of dispersion. Figures at the bottom (**C**, **F**) show the mineralization dynamics predicted in the heterogeneous percolation experiment. Results obtained with an anisotropic dispersion coefficient are indicated as the main lines, while results obtained with an isotropic dispersion are indicated in transparence (pale lines). The ratio of longitudinal dispersion coefficient $d_{disp,\,L}$ over transverse dispersion coefficient $d_{disp,\,T}$ considered in the anisotropic dispersion was equal to 10 as an admissible maximum boundary (Bijeljic and Blunt, 2007). The anisotropic dispersion coefficients were chosen in relation with the effective isotropic dispersion coefficient $d_{disp}$ such as $\frac{1}{3}d_{disp,L} + \frac{2}{3}d_{disp,T} = d_{disp}$. For representation and legend, see **Fig. 3**.

The anisotropy of dispersion slightly shifted the optimal dispersion towards greater values, but this effect was limited. The mineralization dynamics remained similar, and a better fit was even obtained for the best-fitting parameter set under the Contois model.

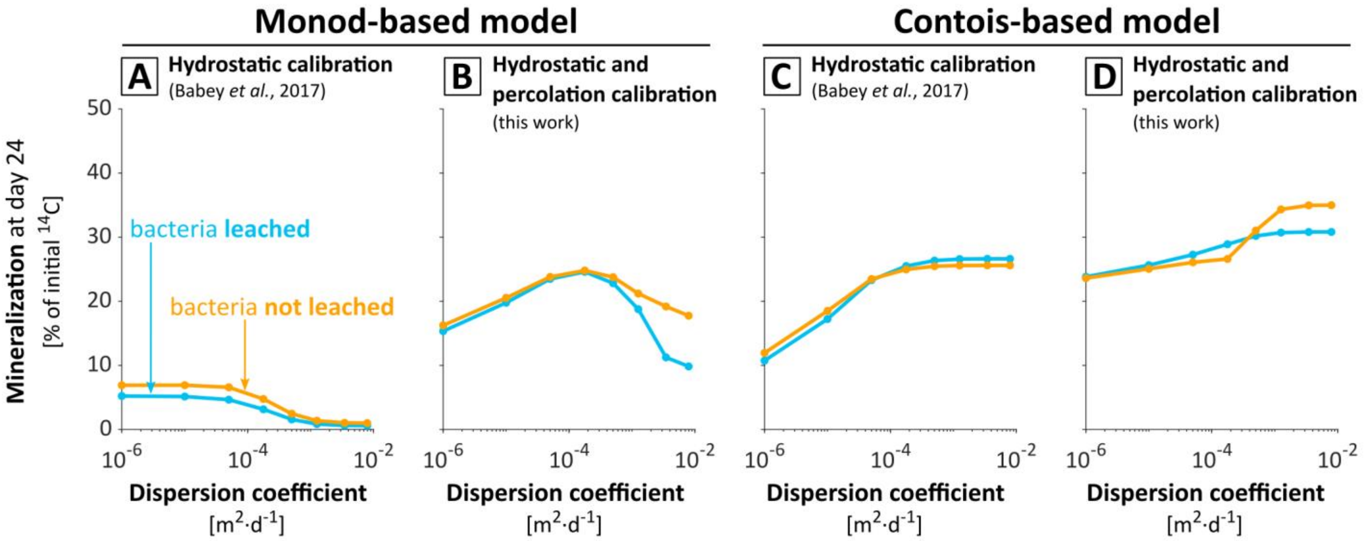


**Fig. S4**. Impact of dispersion on mineralization predicted at day 24, depending on the scenario of bacteria leaching. During the percolation events, bacteria were either leached at the same rate as the substrate (blue lines) or not leached (orange lines). The panels represent the relation between the dispersion coefficient $d_{disp}$ and the final mineralization for the hydrostatic Monod calibration (Babey et al., 2017) applied to the Monod-based (**A**) and the Contois-based (**C**) models, and for the two parameterizations calibrated in this work on both hydrostatic and percolation experiments under Monod-based (**B**) and Contois-based (**D**) models. The parameterizations calibrated under the Monod-based and the Contois-based models are described respectively in sections 3.1 and 3.3, and parameter values are given in **table 2**. As illustrated here, bacteria leaching did not affect the existence of an optimal dispersion for bacteria and its dependence on the maximum uptake efficiency.

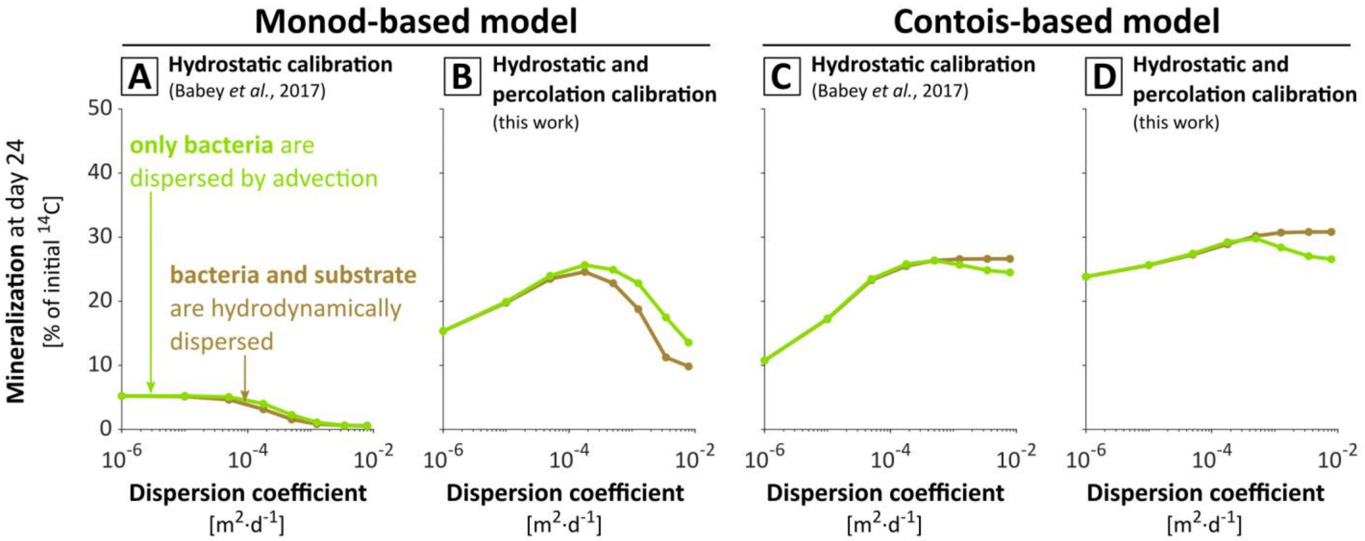


**Fig. S5**. Impact of dispersion on mineralization predicted at day 24, with or without hydrodynamic dispersion of substrate. During the percolation events, either both substrate and bacteria were hydrodynamically dispersed (brown lines) or only bacteria were dispersed (green lines). Panels represent the relation between the dispersion coefficient $d_{disp}$ and the final mineralization for the hydrostatic Monod calibration (Babey et al., 2017) applied to the Monod-based (**A**) and the Contois-based (**C**) models, and for the two parameterizations calibrated in this work on both hydrostatic and percolation experiments under the Monod-based (**B**) and the Contois-based (**D**) models. The parameterizations calibrated under the Monod-based and Contois-based models are described respectively in sections 3.1 and 3.3, and parameter values are given in **table 2**. As illustrated here, hydrodynamic substrate dispersion did not affect the existence of an optimal dispersion for bacteria and its dependence on the maximum uptake efficiency.

### *S.4. Supplementary results*

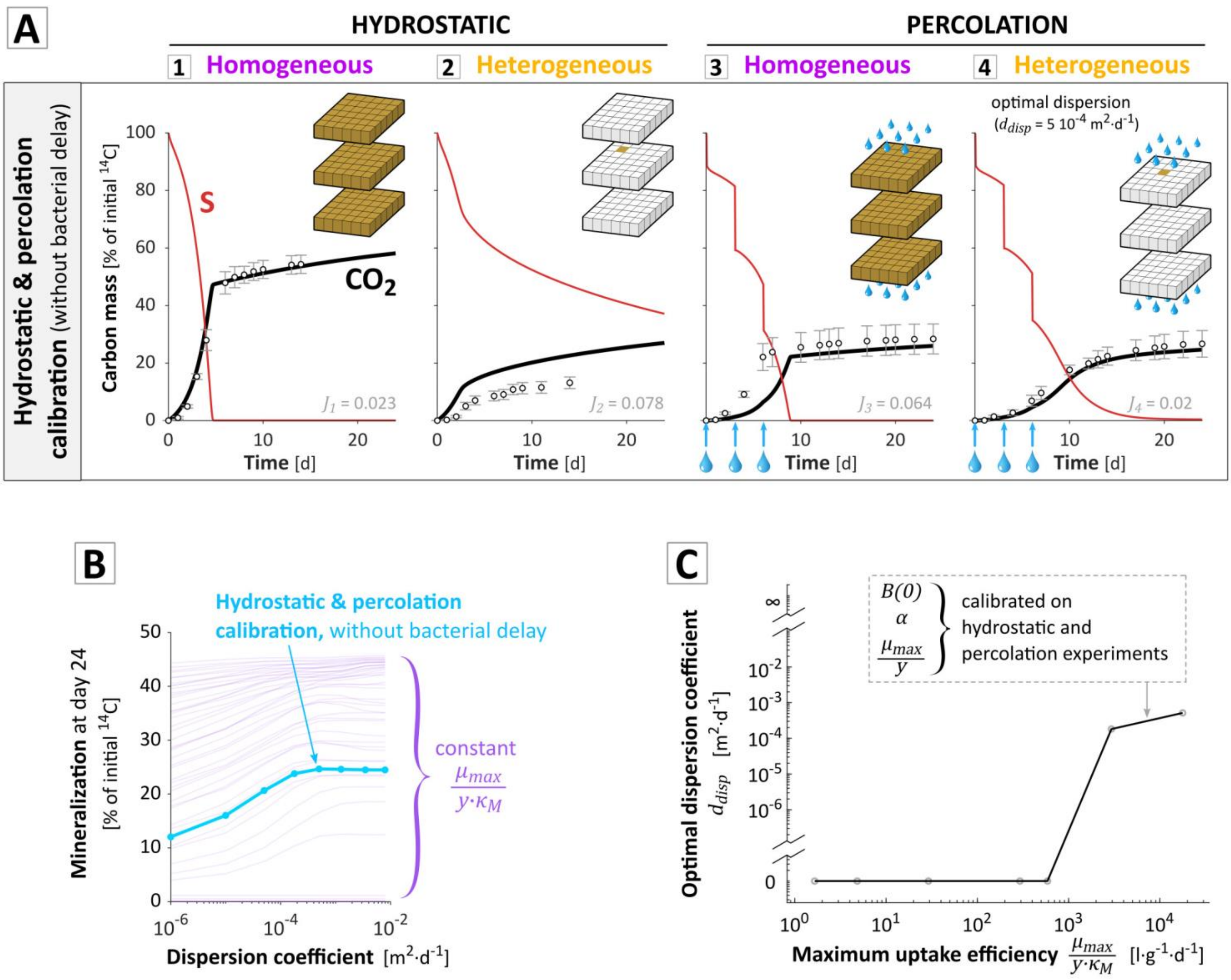


**Fig. S6**. Main modeling results without the explicit bacterial accommodation rate $\alpha$ based on Patarinska et al. (2000). This accommodation function induces a permanent delay in the observed specific growth rate $\mu$, which tends towards its value derived from the Monod relation, following a concave relation whose half-life is given by $\alpha$. For representation and legend, see **Fig. 3**, **4** and **6**.

As discussed in Babey et al. (2017), the inclusion of a delay function in the expression of the bacterial activity was necessary to adequately fit the experimental data of Pinheiro et al. (2015). Nevertheless, the case $\alpha$ = 934.1 $d^{-1}$, which corresponds to a negligible lag phase of 1min30sec, has been included in the screening design. The comparison of those sets with all the sets shows that the parameterization best predicting the data displays a maximum uptake efficiency six times higher, an optimal dispersion coefficient three times higher, and a maximum uptake rate eight times lower. This best-fitting parameter sets predicts well the hydrostatic homogeneous and percolation heterogeneous experiments, but fails to capture the initial dynamics of the two other experiments, for a global discrepancy 1.6 times larger. As shown by the initial dynamics of the mineralization curves, it was possible to reproduce the progressive start of the mineralization without any delay parameter, due to the multiplication of a small initial bacterial population, but it became impossible to find a parameter set in adequacy with all the experimental data.

In regards to the model complexity, it is important to note that, aside from the fact that sorption, bacterial delay and nutrient recycling processes were necessary to adequately fit the experimental data, they did not interfere with the numerical justifications of the following concepts:

- Existence of an optimal bacterial dispersion resulting from a balance between substrate dilution and bacterial density
- Dependence of the optimal dispersion coefficient upon the maximum uptake efficiency
- Significance of the regulation of mineralization by density under low bacterial population densities and high maximum uptake efficiencies commonly found in bulk soil

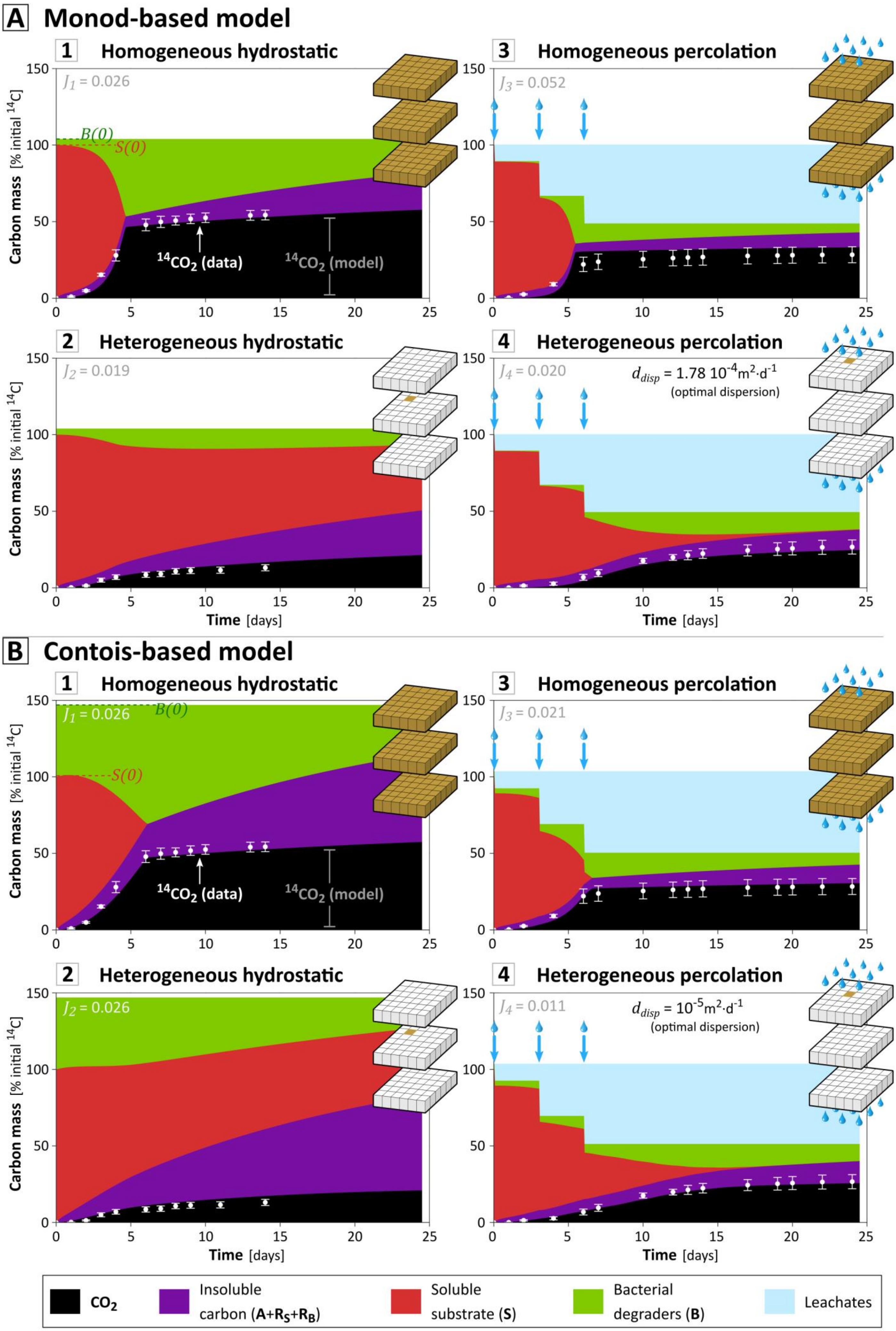
A Monod-based model
1 Homogeneous hydrostatic
J1 = 0.026
B(0)
S(0)
14CO2 (data)
14CO2 (model)
2 Heterogeneous hydrostatic
J2 = 0.019
3 Homogeneous percolation
J3 = 0.052
4 Heterogeneous percolation
J4 = 0.020
ddisp = 1.78 10-4m2·d-1
(optimal dispersion)
B Contois-based model
1 Homogeneous hydrostatic
J1 = 0.026
B(0)
S(0)
14CO2 (data)
14CO2 (model)
2 Heterogeneous hydrostatic
J2 = 0.026
3 Homogeneous percolation
J3 = 0.021
4 Heterogeneous percolation
J4 = 0.011
ddisp = 10-5m2·d-1
(optimal dispersion)
Carbon mass [% initial 14C]
Time [days]
CO2
Insoluble carbon (A+RS+RB)
Soluble substrate (S)
Bacterial degraders (B)
Leachates

**Fig. S7**. Carbon mass balance for the Monod-based (**A**) and the Contois-based (**B**) models calibrated on both hydrostatic and percolation experiments. The different experiments are recalled above each of the graphs and sketched next to them according to **Fig. 1C**. Colors represent the repartition of carbon mass within the four pools described in **Fig. 2** over the whole soil column, normalized by the initial carbon mass of substrate. The initial substrate $^{14}$C-2,4-D (red curve, $S$) is eventually mineralized to $^{14}CO_2$ (black curve), incorporated by the bacterial degraders into their biomass (green curve, $B$) or transferred to insoluble pools (purple curve) as residues sorbed to the soil particles or residues from dead $^{14}$C-bacterial degraders. When the soluble substrate vanishes after some days in [1], [3] and [4], mineralization still occurs but at a much slower rate as it is fueled by nutrient recycling, a feature well captured by the model (Babey et al., 2017). The blue curve refer to the carbon leached at each percolation event (blue arrows) from the soluble substrate and the bacteria. Experimental values and standard deviations of $^{14}CO_2$ production are represented in white. Agreement between experiments and model can be visually assessed by the proximity of the white line to the black area, and the discrepancy $J_q$ between experimental and simulated $^{14}CO_2$ values for each experiment (Eq. (15)) is displayed on top of each graph.

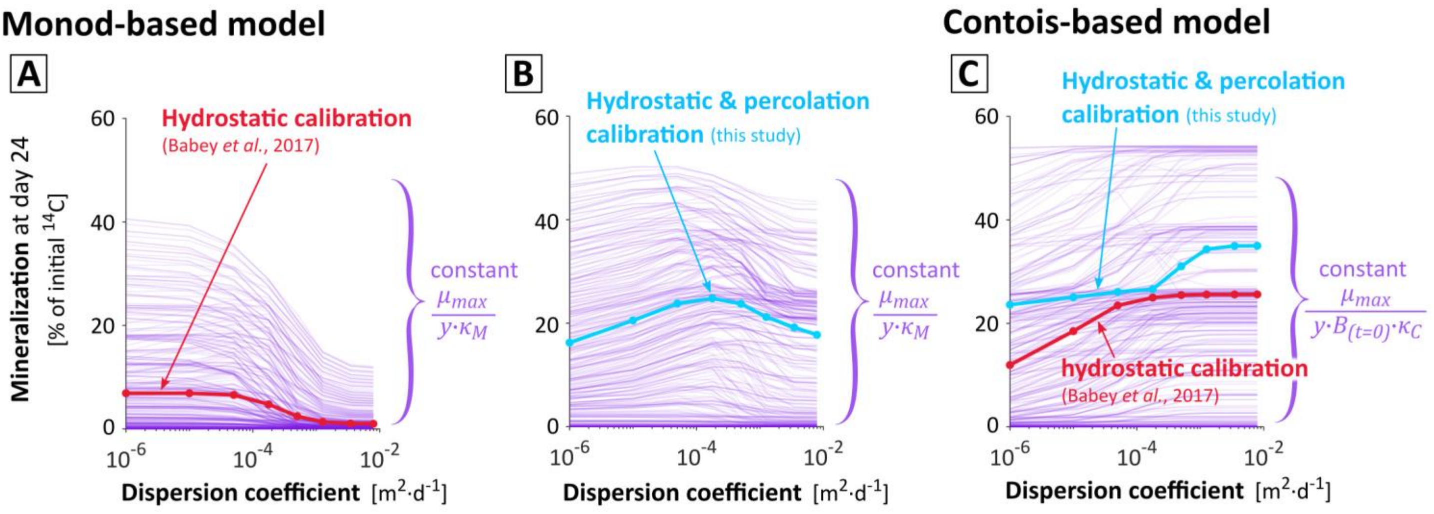


**Fig. S8**. Influence of the dispersion coefficient $d_{disp}$ on mineralization predicted at day 24 $m_{CO2}(t=24)$ for the biological parameter sets calibrated on the sole hydrostatic experiments under the Monod model (**A**, thick red line) and the Contois model (**C**, thick red line), and calibrated on both hydrostatic and percolation experiments under the Monod model (**B**, thick blue line) and the Contois model (**C**, thick blue line). The thin mauves lines correspond to parameter sets with the same specific maximum uptake efficiency, either $(1/y)\cdot\mu_{max}/\kappa_M$ or $(1/y)\cdot\mu_{max}/\kappa_C B(t=0)$ for Monod or Contois model respectively, but different maximum specific uptake rate $(1/y)\cdot\mu_{max}$, accommodation rate $\alpha$ and initial bacterial population density $B(t=0)$. The maximum uptake efficiency is the main determinant of the dispersion leading to the highest final mineralization (see section 3.2.3) while the other biological parameters determine the corresponding mineralization level.

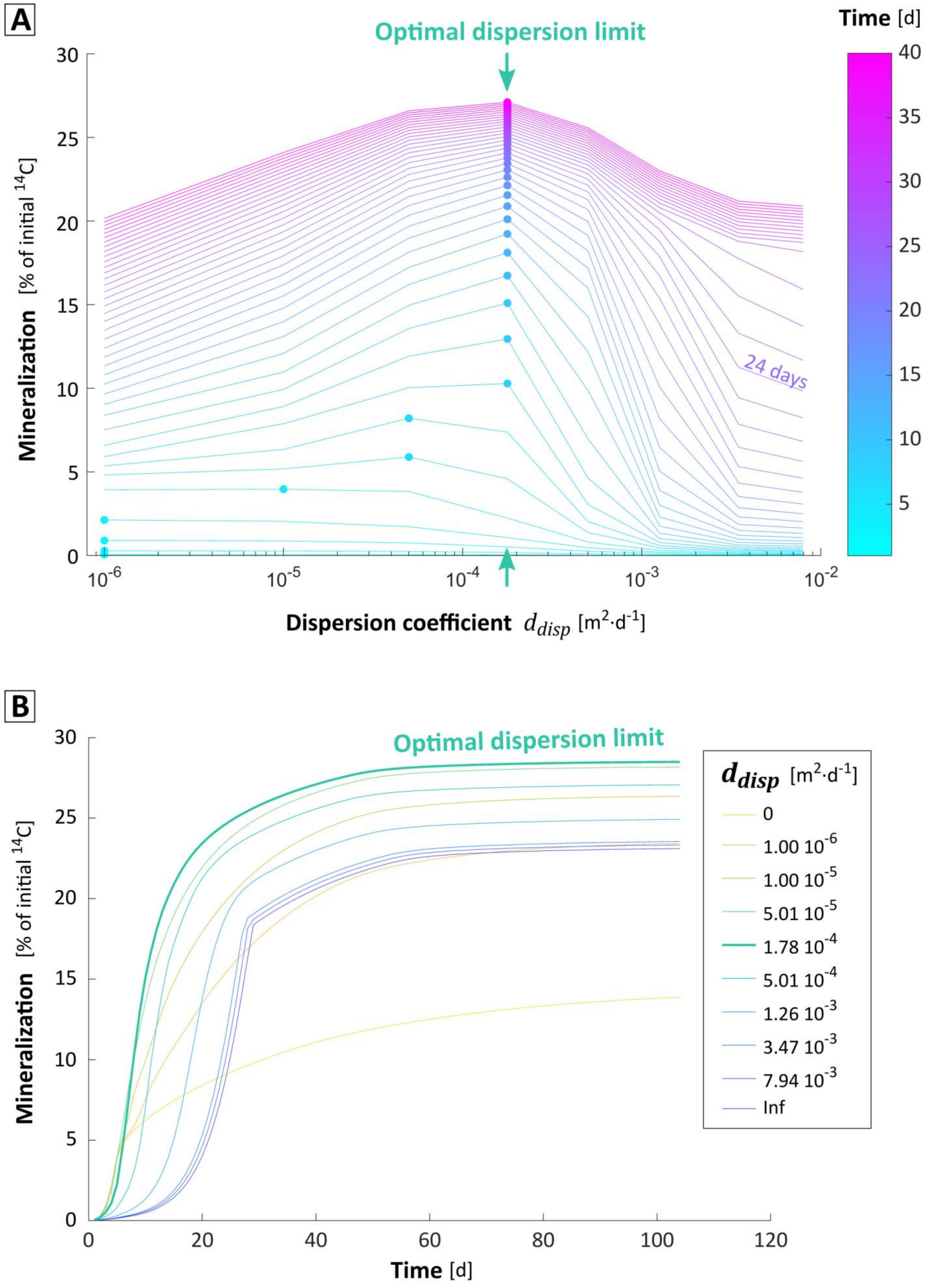

A
Optimal dispersion limit
Time [d]
Mineralization [% of initial 14C]
24 days
Dispersion coefficient d_disp [m2·d-1]
B
Optimal dispersion limit
d_disp [m2·d-1]
0
1.00 10-6
1.00 10-5
5.01 10-5
1.78 10-4
5.01 10-4
1.26 10-3
3.47 10-3
7.94 10-3
Inf
Mineralization [% of initial 14C]
Time [d]

**Fig. S9**. Convergence of optimal bacterial spreading with time. **(A)** Daily mineralization values depending on bacteria dispersion coefficients $d_{disp}$, for the parameter set calibrated under Monod model on both hydrostatic and percolation experiments. For each day represented by the color scale, the optimal dispersion is indicated with a dot. **(B)** Dynamics of mineralization for each $d_{disp}$. The bacterial spreading for maximum degradation varies with time as spatial distributions of bacteria and substrate are dynamic. Diffusion and dispersion steadily smoothen heterogeneities, while bacteria growth and consumption tend to increase them. During the first days, increasing dispersion decreases degradation initial velocity but makes it possible to eventually reach a higher mineralization by delaying the onset of competition for substrate. Nevertheless, this effect vanishes after 4-7 days and increasing dispersion only decreases mineralization. The optimal bacterial dispersion thus tends towards a finite limit of 1.78 $10^{-4}$ $m^2 \cdot d^{-1}$. The optimal dispersion converges as time increases because bacterial growth and consumption are limited in the analyzed experiment by the initial substrate quantity, and because of persistent sorption, carbon stabilization and bacterial decaying favoring quick kinetics.

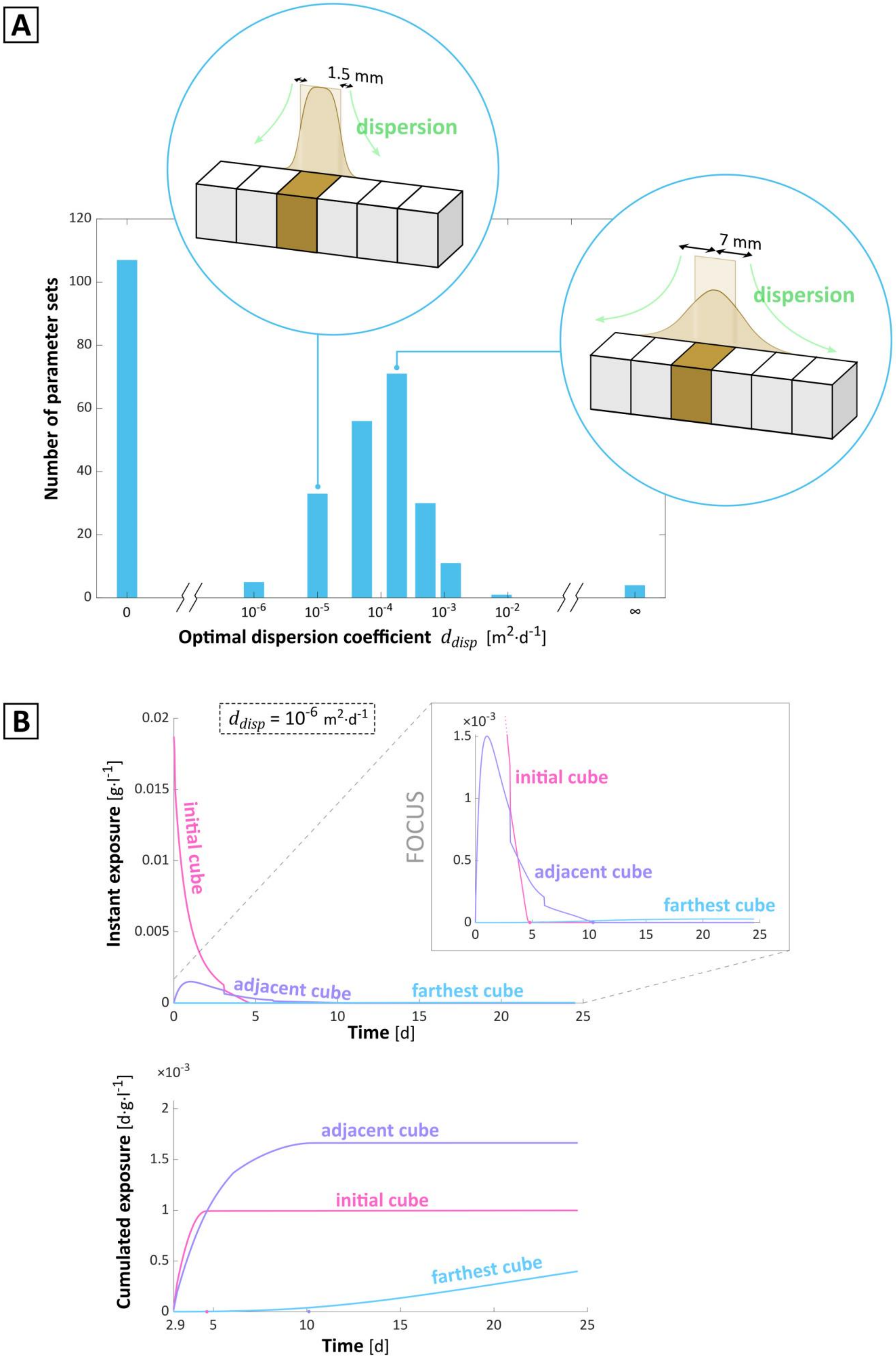
A
1.5 mm
dispersion
7 mm
dispersion
Number of parameter sets
120
100
80
60
40
20
0
0
$10^{-6}$
$10^{-5}$
$10^{-4}$
$10^{-3}$
$10^{-2}$
∞
Optimal dispersion coefficient $d_{disp}$ [$m^2 \cdot d^{-1}$]
B
$d_{disp} = 10^{-6}\ m^2 \cdot d^{-1}$
Instant exposure [$g \cdot l^{-1}$]
0.02
0.015
0.01
0.005
initial cube
adjacent cube
farthest cube
Time [d]
$\times 10^{-3}$
FOCUS
initial cube
adjacent cube
farthest cube
Cumulated exposure [$d \cdot g \cdot l^{-1}$]
$\times 10^{-3}$
adjacent cube
initial cube
farthest cube
2.9
Time [d]

**Fig. S10**. Relevance of bacterial spreading. **(A)** Bar-plot distribution of dispersion coefficients leading to the highest mineralization. The parameter sets considered are the 318 biological parameter sets leading to the best adequacy with data on all experiments except the heterogeneous percolation experiment ($J_{123} < 0.1$). For each of these parameter sets, the dispersion coefficient $d_{disp}$ leading to the best adequacy with data on the percolation experiment is determined, and its distribution is bar-plotted in this graph. Apart from the absence of dispersion ($d_{disp} = 0$), optimal dispersion coefficients roughly follow a lognormal distribution, showing that extremely low and high dispersions are suboptimal. The large number of models without dispersion ($d_{disp} = 0$) likely accounts for all the potential bacterial spreading values equal and inferior to the initial bacterial spreading (one cube) and not individually tested in the screening. Yet, more than 45% of the optimal dispersion coefficients are lower or equal to $10^{-5}$ $m^2 \cdot d^{-1}$, which corresponds to a root-mean-square displacement less than 1 mm, highlighting that bacterial significant spreading is detrimental for half of the screened biological parameter sets. Bacterial spreading can strongly decrease the performance even in our semi-closed system without any loss of substrate by diffusion in the environment. **(B)** Simulated instant and cumulated exposure to substrate concentration in the cube initially amended with substrate (mauve lines), in one of its adjacent cubes (pink lines) and in a cube 36 mm away (blue lines), under the Monod-based model with the biological parameters calibrated on both hydrostatic and percolation experiments and a low dispersion ($d_{disp} = 10^{-6}$ $m^2 \cdot d^{-1}$). Concentration exposures are expressed in mass of 2,4-D per volume of water. The substrate concentration in the initial cube reaches zero after 4.8 days, and after 10 days in the adjacent cube. In addition, due to accommodation time ($1/\alpha = 11$ d) and due to the time necessary for bacterial growth, degradation is small at early times. 95% of the final mineralization occurs after 2 d and 21 h. Therefore, cumulated exposure is represented starting from this date ($t = 2.75$ d), and shows that, for this high-degradation period, the location with the highest cumulated exposure is the adjacent cube.

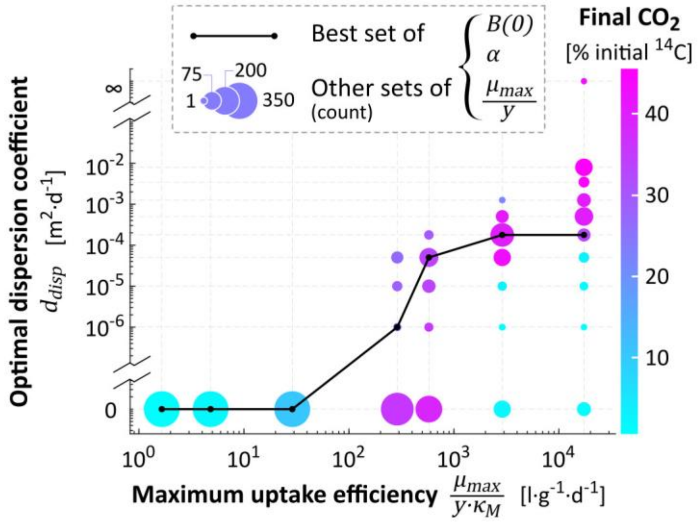


**Fig. S11**. Impact of maximum uptake efficiency on the dispersion coefficient giving the highest mineralization at day 24 for the heterogeneous percolation experiment. As illustrated for the calibrated biological parameterization (black line), the optimal dispersion increases with the maximum uptake efficiency. It is a general feature to all parameterizations. Color scale visually assesses the level of mineralization. With maximum uptake efficiencies below or equal to 28.9 l·g$^{-1}$·d$^{-1}$, mineralization is low no matter the dispersion, even with the highest maximum specific uptake rate $(1/y)\cdot\mu_{max}$, the highest initial biomass $B(t{=}0)$ and without any accommodation delay $\alpha$. After 24 days, it still remains at least 27% of the non-leached initial solute $S(t{=}0)$ in the soil column, versus less than 1% with a maximum uptake efficiency ten times higher. Indeed, with maximum uptake efficiencies of 289 l·g$^{-1}$·d$^{-1}$ or higher, mineralization can reach much higher values. Pink dots highlight that mineralization tend to increase when the maximum uptake efficiency and the corresponding optimal dispersion coefficient both increase.

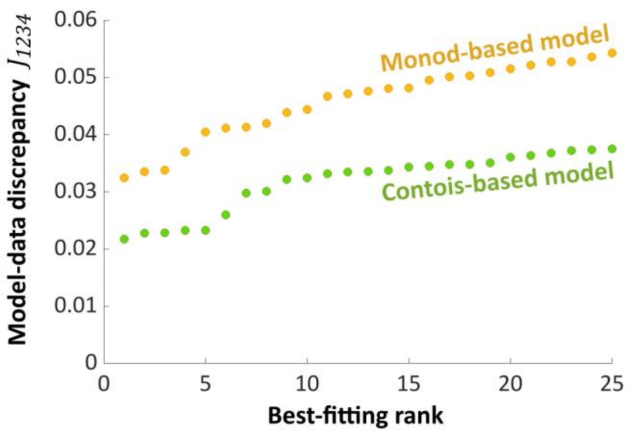


**Fig. S12**. Overall discrepancy $J_{1234}$ (Eq. (15)) between simulations and experiments for the 25 parameter sets leading to the best adequacy with data, under Monod-based model (substrate-dependence, in orange) and Contois-based model (ratio-dependence, in green).

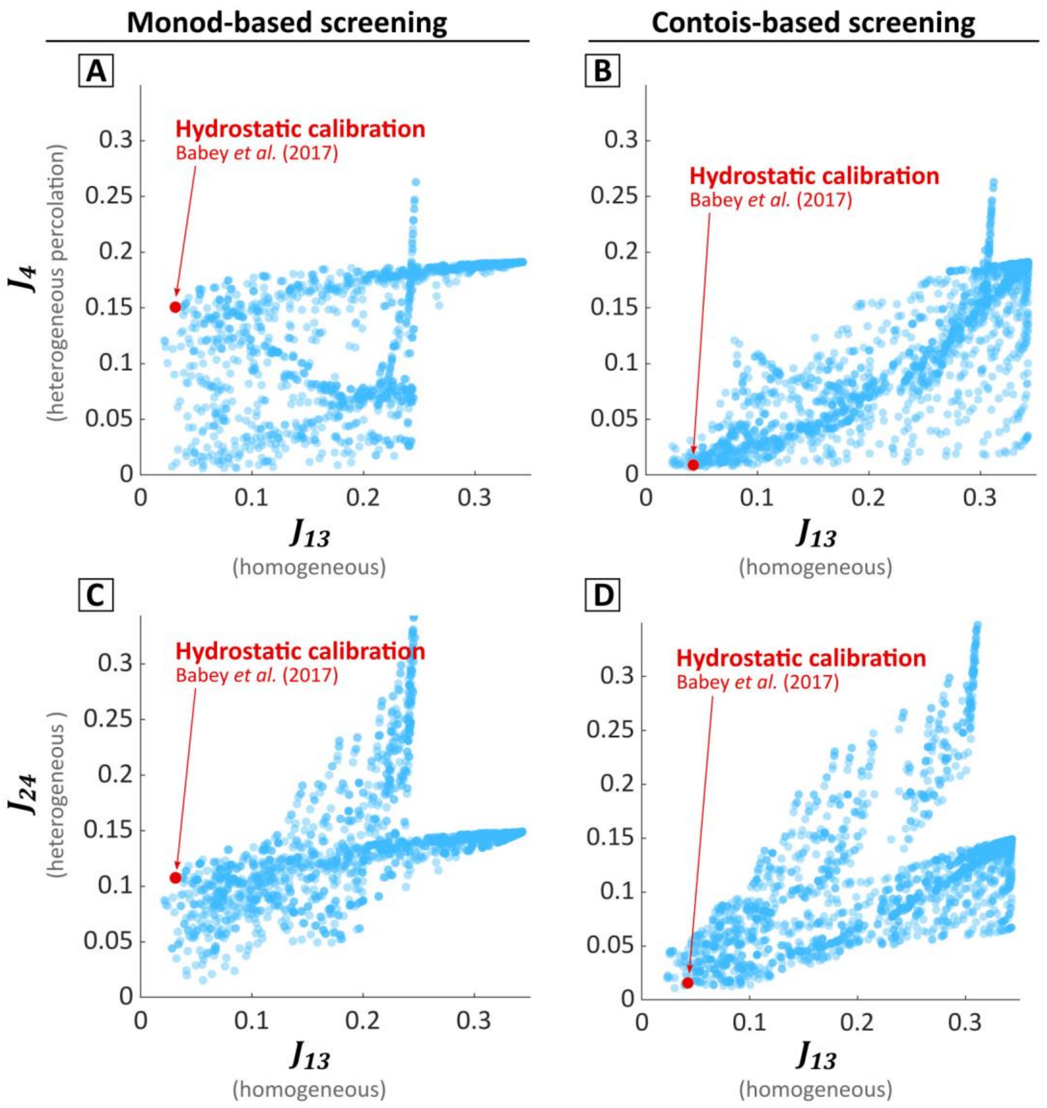


**Fig. S13**. Predictive capacities of Monod-based (**A, C**) and Contois-based (**B, D**) models for aggregated and dispersive conditions. Each dot refers to a set of biological parameters. $d_{disp}$ is chosen for each of the biological parameter set as the dispersion coefficient giving the lowest discrepancy with data. The red dot represents the set calibrated on the sole hydrostatic experiments. The blue dots refer to the other parameter sets. The large vertical dispersion of blue dots on the figures (**A**) and (**C**), even for X-values approaching zero, shows that, with Monod-based model, a good agreement of the model with the homogeneous experiments does not guarantee predictive capacities for the heterogeneous experiments. Conversely, a good agreement with the heterogeneous experiments does not guarantee a good performance on the other experiments. On the contrary, with the Contois-based model, a good agreement of the model with the homogeneous experiments guarantees better predictive capacities for the heterogeneous experiments.

### *S.5. Theoretical collision frequency between bacteria and solute substrate*

The equation of Smoluchowski (1918) for the aggregation of colloids can be used to calculate the collision frequency between bacteria and their substrate in solution, considering both as spherical particles. The frequency of encounter between one bacterium and its substrate is:

$$-\frac{d\nu_S}{dt} = 4\pi(r_S + r_B)(D_S + D_B)\left[1 + \frac{r_S + r_B}{\sqrt{\pi(D_S + D_B)t}}\right]\nu_S\nu_B$$

where $_S$ and $_B$ indices refer to substrate and bacteria respectively, $\nu$ are the number densities, $D$ the molecular diffusion coefficients in water and $r$ the radii.

For $D_S \gg D_B$, $r_B \gg r_S$ and $t > 10^{-3}s$, the equation simplifies to the equation (11) of Smoluchowski:

$$-\frac{d\nu_S}{dt} = 4\pi r_B D_S \nu_S \nu_B$$

This collision frequency for each bacteria can be converted with a change of units as a time differential equation of substrate concentration per gram of bacteria, as:

$$-\frac{dS}{dt} = 4\pi r_B D_S SB = u_{max}SB$$

where $u_{max}$ is the theoretical collisional limit for the maximum uptake efficiency $(1/y)\cdot\mu_{max}/\kappa_M$ (Koch, 1971; Abbott and Nelsestuen, 1988).

With $D_S$ taken equal to 6.9 $10^{-5}$ $m^2 \cdot d^{-1}$ (Saxena et al., 1974) and $r_B$ estimated at 0.75 µm assuming a bacterial volume-weight ratio of 6.22 $10^6$ $\mu m^3 \cdot \mu g^{-1}$ (Balkwill et al., 1988), $u_{max}$ = 6.50 $10^{-7}$ $l \cdot d^{-1}$ (volume of water per bacterial cell per unit of time), which is 135 times higher than the largest maximum uptake efficiency tested in the screening (4.85 $10^{-9}$ $l \cdot d^{-1}$). With a change of units, estimating $m_B$ the average mass of a single 2,4-D degrader = 2.8 $10^{-13}$ g (Dechesne et al., 2010), the maximum uptake efficiency values is $u_{max}$ = 2.13 $10^4$ $g \cdot \mu g^{-1} \cdot d^{-1}$ (mass of dry soil per mass of bacterial carbon per unit of time) = 2.32 $10^6$ $l \cdot g^{-1} \cdot d^{-1}$ (volume of water per mass of bacteria per unit of time). This theoretical limit is similar to the theoretical value of 7.68 $10^6$ $l \cdot g^{-1} \cdot d^{-1}$ assumed by Button (1993) for a marine ultramicrobacteria.

Note that the normalizing bacterial units (bacterial cell, bacterial mass, bacterial carbon mass) depend finely on surface-volume ratio of bacteria and thus on their diameter.
